# It happened again: convergent evolution of acylglucose specialized metabolism in black nightshade and wild tomato

**DOI:** 10.1101/2021.06.08.447545

**Authors:** Yann-Ru Lou, Thilani M. Anthony, Paul D. Fiesel, Rachel E. Arking, Elizabeth M. Christensen, A. Daniel Jones, Robert L. Last

## Abstract

Plants synthesize myriad phylogenetically-restricted specialized (aka ‘secondary’) metabolites with diverse structures. Metabolism of acylated sugar esters in epidermal glandular secreting trichomes across the Solanaceae (nightshade) family are ideal for investigating the mechanisms of evolutionary metabolic diversification. We developed methods to structurally analyze acylhexose mixtures by 2D NMR, which led to the insight that the Old World species black nightshade (*Solanum nigrum*) accumulates acylglucoses and acylinositols in the same tissue. Detailed *in vitro* biochemistry – cross validated by *in vivo* virus induced gene silencing – revealed two unique features of the four-step acylglucose biosynthetic pathway: a trichome-expressed, neofunctionalized invertase-like enzyme, SnASFF1, converts BAHD-produced acylsucroses to acylglucoses, which in turn are substrates for the first-reported acylglucose acyltransferase, SnAGAT1. This biosynthetic pathway evolved independently from that recently described in the wild tomato *S. pennellii*, reinforcing that acylsugar biosynthesis is evolutionarily dynamic with independent examples of primary metabolic enzyme cooption and additional variation in BAHD acyltransferases.

**Teaser:** Analysis of plant protective surface hair chemistry revealed evolutionary mechanisms leading to metabolic innovation.

## Introduction

Plants synthesize myriad structurally diverse specialized (historically referred to as ‘secondary’) metabolites, which provide humans with medicines, food additives and natural insecticides. In recent years, an increasing number of these compounds were documented to serve ecological functions. Specialized metabolites group into taxonomically-restricted classes, such as glucosinolates in Brassicales and benzoxazinoid alkaloids in Poaceae (*1–4*). Because the full diversity of specialized metabolites is distributed across the Kingdom, the vast majority of plant specialized metabolites are uncharacterized. In contrast to the typically conserved enzymes and pathways of primary metabolism, variation of the lineage-specific specialized metabolic pathways is observed within and across species (*2, 5–7*). In fact, specialized metabolic pathway enzymes have characteristics that promote product diversity. These include promiscuity (seen for example in BAHD (**B**EAT, **A**HCT, **H**CBT, and **D**AT) acyltransferases), gene duplication (e.g. cytochromes P450) and neo- and sub-functionalization of gene expression and enzyme activity.

Glandular-secreted trichome acylsugars are a structurally diverse group of specialized metabolites produced in the nightshade (Solanaceae) family that have developed into an exemplary system for investigating the emergence and diversification of novel metabolites in plants (*8–16*). Their anti-herbivory and anti-microbial activities make them of interest in understanding their ecological roles and as breeding targets for crop protection (*17–20*). Most characterized acylsugars consist of a sucrose, glucose or inositol sugar core esterified at various positions with acyl chains that are often aliphatic, differing in length – typically C2-C12 (*8, 16, 21–29*). A single species in Solanaceae can produce dozens of distinct acylsugars consisting of these simple biosynthetic building blocks (*14, 28*).

Sugar esters with a sucrose core (acylsucroses) are predominant in Solanaceae (*8, 14*). The well-characterized biosynthetic pathway producing cultivated tomato (*Solanum lycopersicum*) acylsucroses involves four trichome-expressed BAHD-family **a**cyl**s**ugar **a**cyl**t**ransferases (ASAT1, ASAT2, ASAT3 and ASAT4) (*9–11*). Evolutionarily-related ASATs were characterized in acylsugar biosynthetic pathways in species across the family, with all but one using sucrose or acylsucroses as acceptor substrate (*12–14, 16*). These enzymes sequentially transfer acyl chains from acyl-Coenzyme A substrates (acyl-CoAs) onto specific positions of the sugar core, with individual ASATs varying in both acyl donor and acyl acceptor substrates. Substrate specificity and promiscuity of these enzymes influence inter- and intra-specific diversification of acyl chain length and acylation position as reported in *Petunia axillaris*, *Salpiglossis sinuata,* and wild tomato species (*12–14*).

While the majority of characterized acylsugars are based on sucrose, acylhexoses have also been described. Specifically, acylglucoses were reported in species within the *Solanum*, *Datura* and *Nicotiana* genera (*21, 27, 29–31*), while acylinositols were characterized from the South American native fruit crop naranjilla (*Solanum quitoense*) and the Central American native orangeberry nightshade (*Solanum lanceolatum*) (*16, 23*). Relatively little is reported about acylhexose biosynthesis, with one published example of a BAHD acylinositol acetyltransferase in *S. quitoense* (*16*), and no reported glucose or acylglucose acyltransferases. In fact, we recently showed that wild tomato *S. pennellii* acylglucoses are produced by acylations of sucrose followed by conversion to acylglucoses by a trichome-expressed invertase-like enzyme, **a**cyl**s**ucrose **f**ructo**f**uranosidase 1 (SpASFF1) (*15*). This enzyme utilizes the acylsucrose substrates with all acyl chains on the pyranose ring (P-type acylsucroses) found in *S. pennellii* but not in cultivated tomato or other wild tomato relatives. Hence, this ASFF1-dependent pathway evolved independently in the lineage of *S. pennellii*. This leaves open the question of how acylglucose biosynthesis proceeds outside the tomato sub-clade of the *Solanum* genus.

To address this problem, we investigated trichome acylsugar specialized metabolism in the predominantly Eurasian black nightshade species, *S. nigrum*. In contrast to the New World species *S. pennellii*, which produces both acylglucoses and acylsucroses, our previous work suggested that the trichomes on *S. nigrum* young leaves, stems and fruits exclusively accumulate acylhexoses (*14*). Moreover, while other characterized species accumulate acylsugars with three or more acyl chains, *S. nigrum* accumulates acylhexoses with as few as two acyl chains.

Understanding the enzymology behind acylhexose biosynthesis requires accurate assessment of acylation sites on sugar cores, for which mass spectrometry yields limited information, but Nuclear Magnetic Resonance (NMR) spectroscopy is well suited. Despite the power of modern methodologies, obtaining NMR-resolved chemical structures requires samples of sufficient abundance and purity. As a result, while we have liquid chromatography–mass spectrometry (LC–MS) data for hundreds of structurally diverse acylsugars, only a subset have unambiguous structure annotations. For example, of more than 38 surveyed Solanaceae species that accumulate acylsugars, fewer than half have NMR structures elucidated for at least one acylsugar (*14, 16, 21–26, 28, 29*).

Here we report the structural characterization of *S. nigrum* acylhexoses – consisting of triacylinositols, diacylglucoses and triacylglucoses – and describe characterization of the *S. nigrum* acylglucose biosynthetic pathway. Our accelerated structure elucidation pipeline provided NMR-resolved structures for *S. nigrum* acylsugars from leaf-surface extracts without purifying individual compounds. A combination of *in vitro* biochemical pathway reconstruction and *in vivo* virus-induced gene silencing (VIGS) validated the acylglucose biosynthetic pathway. This pathway consists of four trichome-expressed enzymes. The two acylsucrose acyltransferases SnASAT1 and SnASAT2 produce diacylsucroses, which serve as substrates for an acylsucrose fructofuranosidase 1 (SnASFF1), which synthesizes diacylglucoses. Triacylglucoses are then produced by SnAGAT1. While it is analogous to the *S. pennellii* acylglucose pathway, the *S. nigrum* pathway evolved independently, with co-option of the neofunctionalized SnASFF1 from a distinct lineage of the invertase gene phylogeny, and evolution of a BAHD acyltransferase that acts on acylglucoses. This work demonstrates the value of leveraging analytical chemistry, phylogenetics, enzymology and knowledge gained from model organisms to facilitate chemical structure assignment and biochemical pathway elucidation in a non-model organism.

## Results

### *S. nigrum* accumulates di- and triacylated acylglucoses and triacylated acylinositols

Understanding the atomic connections of products and metabolic intermediates is critical for biochemical pathway elucidation. NMR-resolved structures of purified compounds are the gold standard for small molecule metabolites, but obtaining these data can be time-consuming. We sought to speed up the process by subjecting total extract or partially purified mixtures to 2D NMR analyses, taking advantage of previously developed acylsugar analysis methods (*28, 29, 32*) (Fig. 1*C*).

**Figure 1.**
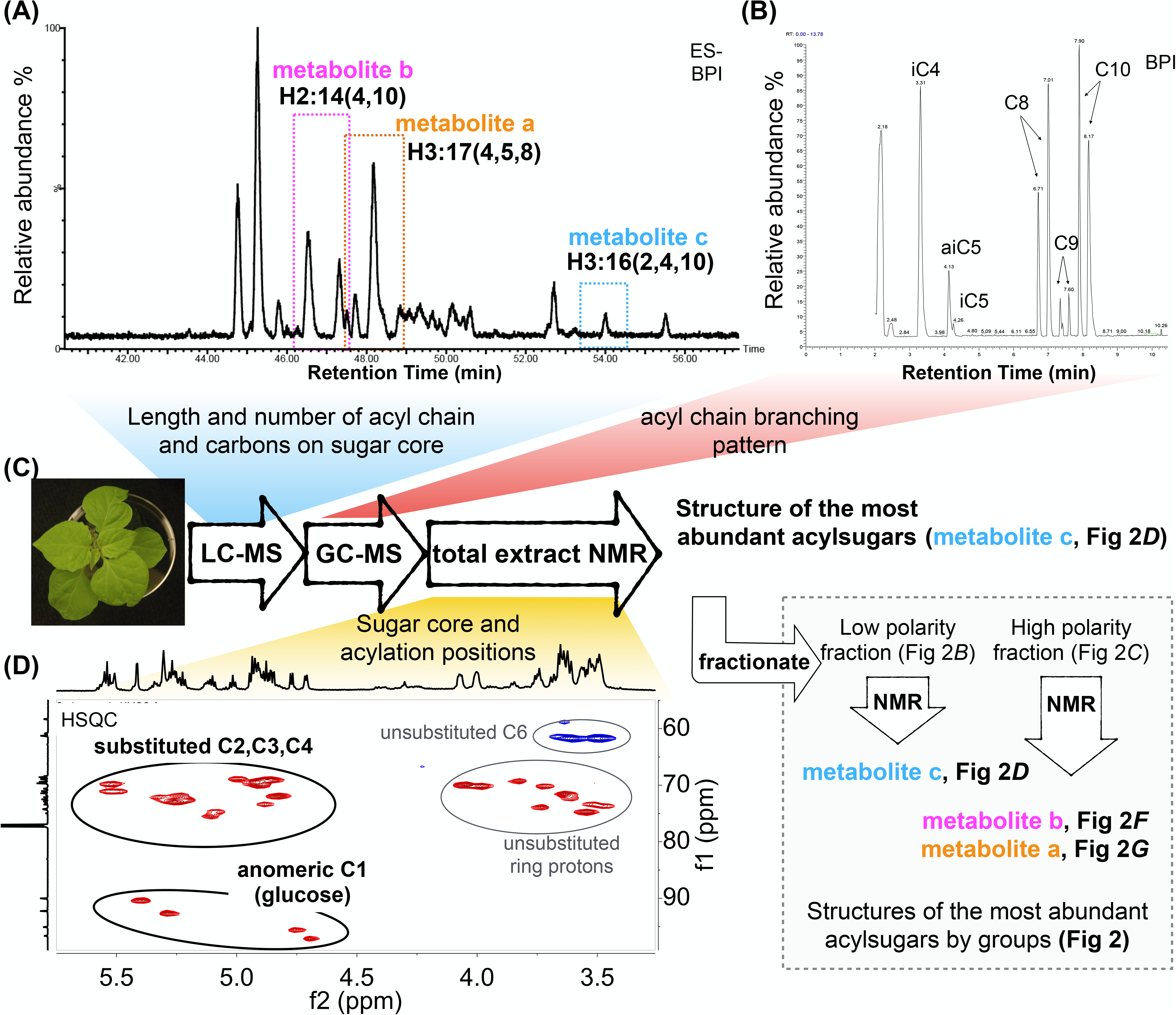
Integrated analysis of *S. nigrum* acylhexose structures. (**A**) LC-MS profile of *S. nigrum* leaf dip extracts reveals acylsugar-like compounds. Metabolites a, b and c represent the most abundant acylinositol, diacylglucose and triacylglucose, respectively. See Fig. S1, S2, S3, Table S1 and Supplementary Methods for details. (**B**) Fatty acid ethyl ester profile of saponified acylsugars analyzed using GC-MS reveal abundant iC4, iC5, aiC5, as well as straight and branched C8, C9 and C10 acyl chains. The abbreviations “i” and “ai” refer to *iso*- and *anteiso*-branched isomers (terminally and sub-terminally branched, respectively). (**C**) Schematic of pipeline used for acylsugar structure determination. (**D**) NMR analysis of total acylsugars revealed that the most abundant acylsugar contains a glucose core and acyl chains on C2, C3 and C4 ring protons. BPI, base peak intensity; ESI-, negative electrospray ionization. All ESI- mode acylsugars are identified as formate adducts. The corresponding *m/z* of these compounds are listed in Table S1 and Supplementary Methods.

As a first step to characterize the number and diversity of acylsugars, we profiled metabolites in *S. nigrum* leaf-dip extracts using LC-MS (Fig. 1*A*). We annotated 99 metabolite features – among them 45 with high confidence based on criteria discussed in the Supplementary Method – with characteristics similar to known acylsugars based on molecular ion adduct masses, mass spectra at elevated collision energies and fractional hydrogen content as quantified by relative mass defect (*32, 33*) (Fig. 1*A*, Fig. S1 and S2, Table S1 and Supplementary Text). In contrast to the majority of Solanaceous plants, we only detected acylhexoses in *S. nigrum* extracts (*14*). LC-MS fragmentation patterns revealed that the acylsugars are typically composed of hexose cores and two or three short-(C2, C4, C5) or medium-chain length (C8, C9, and C10) acyl esters (Table S1 and Supplementary Text). Thirty-nine of the 45 high-confidence putative acylsugars showed fragmentation patterns similar to previously analyzed *S. pennellii* acylglucoses (*29*). However, we observed six triacylhexoses with unusually low fragment ion signals in negative mode (Fig. S2); this is a characteristic reminiscent of *S. quitoense* acylinositols (*16, 34*).

The acylglucose- and acylinositol-like peaks in *S. nigrum* appear as groups of two or more chromatographic peaks sharing identical molecular masses and indistinguishable mass spectra at high collision energy as consistent with multiple isomeric forms (Fig. S3). Extracted ion chromatograms for fragment ion masses characteristic of acyl groups (carboxylate anions) revealed that many individual chromatographic peaks result from multiple co-eluting metabolite features, some of which present indistinguishable mass spectra. For example, the second-eluting peak annotated as H3:16 (Fig. S3) exhibits asymmetry suggestive of four isomers, two of which overlap in chromatographic retention. All four components share the same acyl chain complements – C2, C4 and C10 acyl chains on a hexose core as evidenced from the extracted fragment ion chromatograms that confirm C4 and C10 acyl groups (Fig. S3). We expect each acylglucose to resolve as two peaks corresponding to α and β anomers that interconvert, accounting for the presence of two peaks for each (*29*). The observation of additional peaks led to the hypothesis that four isomeric forms reflected different acyl chain branching patterns rather than differences in acyl group lengths or positions of esterification. This is supported by the ethyl ester profile revealed by gas chromatography-MS (GC-MS) after saponification and transesterification (Fig. 1*B*). The ethyl ester profile demonstrated both abundant straight (nC8, nC9, nC10) and branched (iC4, iC5, aiC5, branched C8, branched C9, branched C10) aliphatic chains (Fig. 1*B*), supporting the conclusion that acyl chain branching variation contributes to the number of structural isomers.

The combined information gained from LC-MS and GC-MS led us to conclude that *S. nigrum* accumulates at least 45 different types of di- and triacylhexoses including structural isomers with acyl chains differing in branching patterns. While unequivocal structural information of these acylhexoses is essential for a complete understanding of their biosynthesis, the chromatography-based analytical techniques do not provide needed resolution or selectivity. For example, MS has not yet developed technologies that identify acylation positions or differentiate isomeric hexose cores (e.g. glucose or inositol). Such information is central to understanding functions of key metabolic enzymes that shape acylsugar chemical diversity. We employed NMR spectroscopy, which provides important information about molecular topology, to obtain these structural details.

We sought to avoid the time-consuming process of purifying metabolites to homogeneity by analyzing unfractionated and partially purified fractions (Fig. 1*C*). This took advantage of the 2D NMR spectroscopy methods Heteronuclear Single Quantum Coherence (HSQC), which provides information about hydrogen-carbon attachments, and Heteronuclear Multiple Bond Correlation (HMBC), which yields information about atoms separated by 2-3 bonds. Analysis of the total, unfractionated acylsugar extracts from young plants revealed the dominant presence of a glucose backbone with acyl chains at the 2-, 3- and 4-positions (Fig. 1*D*). However, abundant triacylglucoses obscured the signals of diacylglucoses and acylinositols in the total extract: we employed partial purification to obtain similar information about these less abundant acylsugars.

Silica gel chromatography was used to separate triacylglucoses (fraction 1) from diacylglucoses and triacylinositols (fraction 2) based largely on the number of hydroxyl groups: there are two hydroxyls on triacylglucoses; three on diacylglucoses and triacylinositols (Fig. 2*A-C*). Analysis of the NMR spectra from fraction 1 confirmed that it predominantly contains triacylglucoses with one medium-length (8-10 carbon) acyl chain esterified to the 4-position, a short (4-5 carbon) acyl chain at the 3-position, and an acetyl (C2) group at the 2-position (Fig. 2*D* and Table S2). In agreement with LC-MS data, NMR spectra of fraction 2 exhibited signals of two distinct groups of acylhexoses in the sugar region (δ_H_ = 3-6 ppm; δ_C_ = 60-100 ppm) (Fig. S4). The anomeric positions in hexoses stand out in having larger (downfield) ^13^C chemical shifts (δ_C_ ∼ 90-100 ppm) than other acylsugar carbon atoms (Fig. S4). We employed 2D HSQC-TOtal Correlation SpectroscopY (HSQC-TOCSY), which generates spectra at long mixing times to provide connectivity information between coupled nuclei in a molecule (*35*). Such information aids in associating sugar core hydrogen and carbon signals within a molecule and distinguishing signals from other molecules in a mixture. This approach provided a useful strategy for distinguishing acylation positions on acylglucoses and acylinositols in fraction 2 where signals often overlap. Our analysis revealed acylations at the 3- and 4-positions on a sugar core with an anomeric center (glucose) and acylations at 2-, 3-, and 4-positions on a hexose core with no anomeric center (inositol) (Fig. 2*E* and Fig. S4). HMBC data analysis confirmed the presence of abundant diacylglucoses with a medium-length acyl chain as R_4_ and a short acyl chain as R_3_ (Fig. 2*F* and Table S3). This approach also revealed the presence of triacylinositols with two short- and one medium-length acyl chain as R_2_, R_3_ and R_4_ (Fig. 2*G* and Table S4).

**Figure 2.**
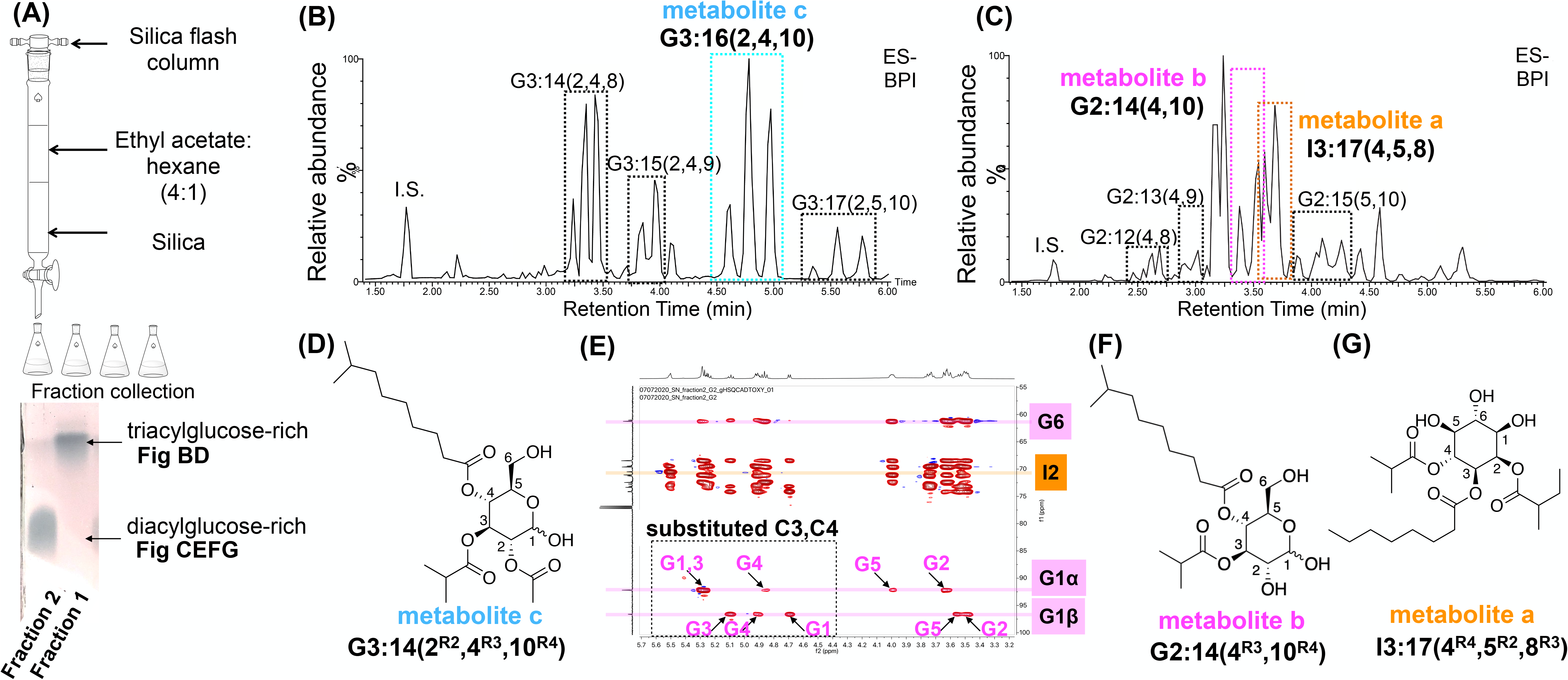
*S. nigrum* accumulates triacylinositols and di- and triacylated acylglucoses. (**A**) Silica flash column separation of leaf dip extracts into less polar fraction 1 and more polar fraction 2, as analyzed by thin layer chromatography. (**B**) LC-MS profile of fraction 1 reveals mostly triacylglucoses. The peaks of metabolite c (cyan dashed rectangle) with identical *m/z* and mass spectra represent the two anomers each of two structural isomers of triacylglucose G3:16(2,4,10). (**C**) LC-MS profile of fraction 2 reveals abundant diacylglucoses and triacylinositols. The two peaks with *m/z* and mass spectra consistent with metabolite a (orange dashed rectangle) represent two structural isomers of I3(4,4,8), whereas metabolite b (magenta dashed rectangle) resolves into multiple peaks representing the two anomers of the two structural isomers of G2:14(4,10). (**D**) NMR-resolved structure of metabolite c, the most abundant triacylglucose in fraction 1. See Table S2 for details. (**E**) HSQC-TOCSY separated spin systems between diacylglucoses and triacylinositols in fraction 2. See Fig. S4 for details. (**F**) NMR-resolved structure of metabolite b, the most abundant diacylglucose, and (**G**) metabolite a, the most abundant triacylinositol in fraction 2. See Table S3 and S4 for details. BPI, base peak intensity; ESI-, negative electrospray ionization. All chromatograms showing telmisartan as internal standard (I.S.). All ESI- mode acylsugars are identified as formate adducts. The corresponding *m/z’s* of these compounds are listed in Table S1.

Analysis of the combined LC-MS, GC-MS and NMR results led to the conclusion that *S. nigrum* accumulates diacylglucoses esterified at the 3- and 4-positions, with triacylglucoses and triacylinositols esterified at the 2-, 3-, and 4- positions. For the remainder of this report we follow the nomenclature established in Schilmiller et al. (2010) to describe these compounds: SX:Y(A^a^,B^b^,C^c^), in which **S** indicates the type of sugar core (**G** for **g**lucose, **I** for **i**nositol), **X** is the total number of substituting acyl chains, **Y** signifies the sum of acyl chain carbons and **(A^a^,B^b^,C^c^)** represents the number of carbons of each chains with the superscript documenting the site of substitution. Based on this nomenclature, G2:14(4^R3^,10^R4^), G3:16(2^R2^,4^R3^,10^R4^), and I3:17(4^R4^,5^R2^,8^R3^) represent some of the most abundant acylsugars in each class (Fig 2 D, F and G). In cases where the identity of the hexose core remains ambiguous, the letter **H** for **h**exose is employed.

### Identification of a trichome-expressed β-fructofuranosidase involved in *S. nigrum* acylsugar biosynthesis

The observation that acylglucoses are predominant in *S. nigrum* trichome extracts led us to posit that, as previously shown for *S. pennellii* LA0716 acylglucose biosynthesis, acylsucroses are intermediates in acylglucose production. To test the hypothesis, we identified trichome-expressed transcripts in *S. nigrum* predicted to encode proteins of the glycoside hydrolase family 32, which includes SpASFF1. Twenty invertase-like homologs were identified by BLAST from *S. nigrum* RNAseq data (Fig. S5*A*). We reasoned that – as acylsugars appear to accumulate on *S. nigrum* trichome tip cells (Fig. S5*B*) – the acylglucose biosynthetic genes in *S. nigrum* would be highly expressed and enriched in trichomes. Among the twenty homologs, six demonstrate trichome-enriched expression (Fig. S5*C*). Five of the six have sequence lengths comparable to functional invertases, and have WXNDPNG, RDP and EC sequences characteristic of β-fructofuranosidases (Fig. S5*D*). Because we previously observed a **P**ST**P** non-canonical substrate binding site in SpASFF1(*15*), we sought invertase-like sequences also lacking the canonical DXXK in *S. nigrum* trichome expressed transcripts. Indeed, assembly c70979_g1 has **P**LT**Y** in place of that conserved substrate binding site (Fig. S5*D*). Based on the *in vivo* transient silencing and *in vitro* enzyme activities results described below, we designated this protein SnASFF1.

We developed and deployed *S. nigrum* VIGS to test the hypothesis that SnASFF1 is involved in *S. nigrum* acylglucose biosynthesis *in vivo*. Plants with silenced *SnASFF1* accumulated diacylsucroses (referred to as ‘*in vivo* diacylsucroses’ below) that were not detected in the control plants (Fig. 3, Fig. S5*E* and *F* and Table S5). NMR spectra of the early- and late-eluting S2:14(4, 10) isomers revealed that the acylation positions – on positions 3 and 4 (Fig. S5*G*) – mirror those of *S. nigrum* acylglucoses (Fig. 2*D* and *F*), supporting their role as intermediates in the acylglucose biosynthetic pathway. As characterized ASATs show acylation position specificity, this result is consistent with the hypothesis that there are two or more BAHD acyltransferases that produce diacylsucrose substrates for this SnASFF1 invertase-like enzyme.

**Figure 3.**
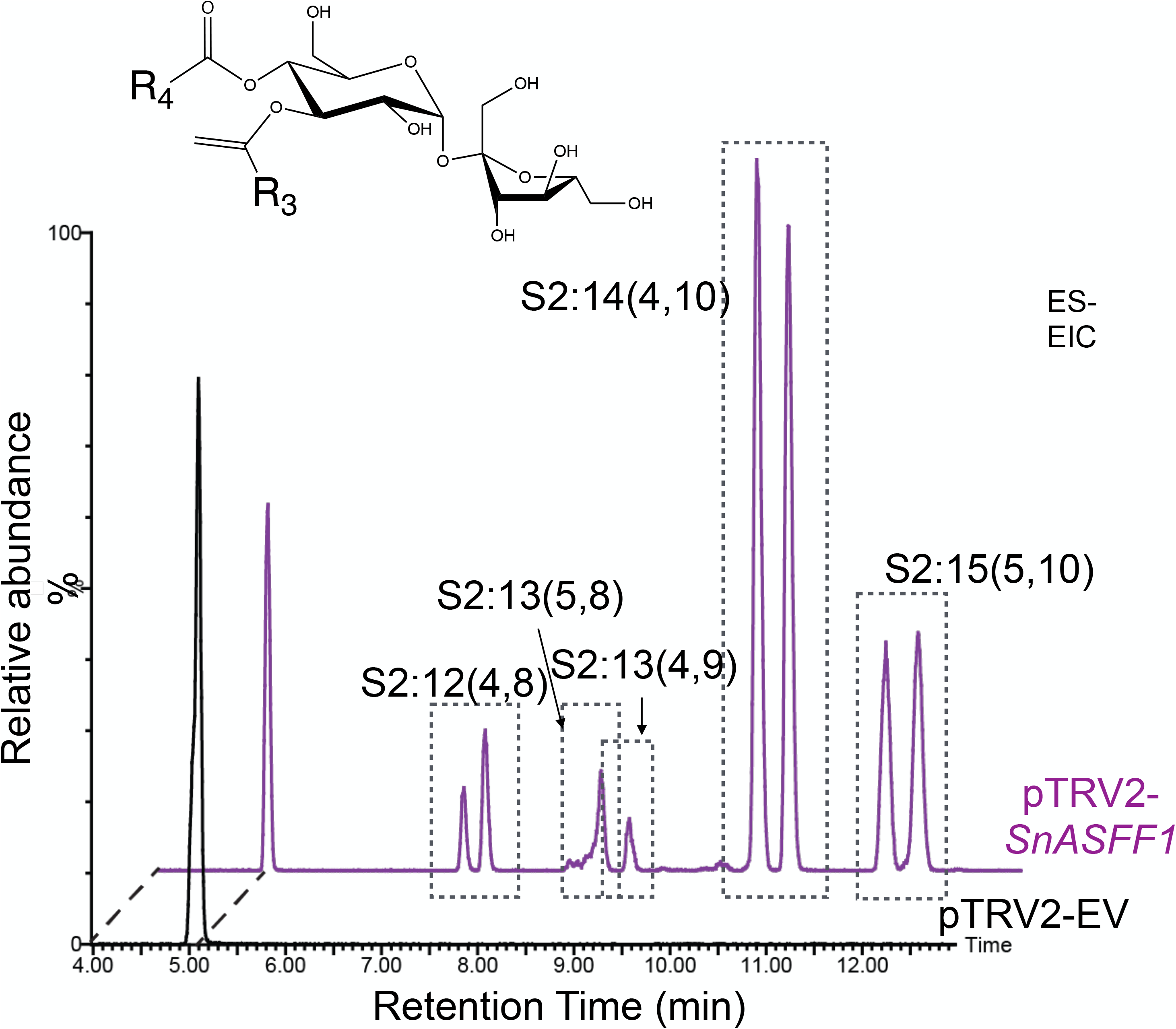
Diacylsucroses accumulate in *SnASFF1* virus induced gene silencing (VIGS) lines. LC-MS analysis of acylsugars extracted from *SnASFF1*-targeted (purple trace) and empty vector VIGS plants (black trace) show that *SnASFF1*-silenced lines accumulate diacylsucroses that are undetectable in control plants. The acylation at positions 3 and 4 on S2:14(4,10) was verified by NMR as shown in Fig. S5. Extracted ion chromatogram (EIC) values indicate telmisartan as internal standard (I.S.) [*m/z* 513.23] and the formic adducts of S2:12 (*m/z* 583.26), S2:13 (*m/z* 597.28), S2:14 (*m/z* 611.29), and S2:15 (*m/z* 625.31). ESI-, negative electrospray ionization.

### Identification of trichome-specific BAHD acyltransferases

Based on published results with acylsucrose- and acylinositol-producing plants in Solanaceae, we hypothesized that acyltransferase enzymes of *S. nigrum* acylsugar biosynthesis would be trichome-enriched and evolutionarily related to characterized ASATs (*14, 16*). The combined tissue-specific expression and homology-guided approach led us to consider six highly trichome-enriched BAHD family members in the *S. nigrum* transcriptome (Fig. 4*A* and Fig. S6). These candidates are predicted to encode proteins with characteristics of active BAHD family enzymes (*36*). First, they have the conserved HXXXD catalytic motif and the DFGWG-like structural motif (Fig. S7). Second, the predicted proteins all have lengths comparable to characterized functional BAHDs (*8–16*) (Fig. S6*B*).

**Figure 4.**
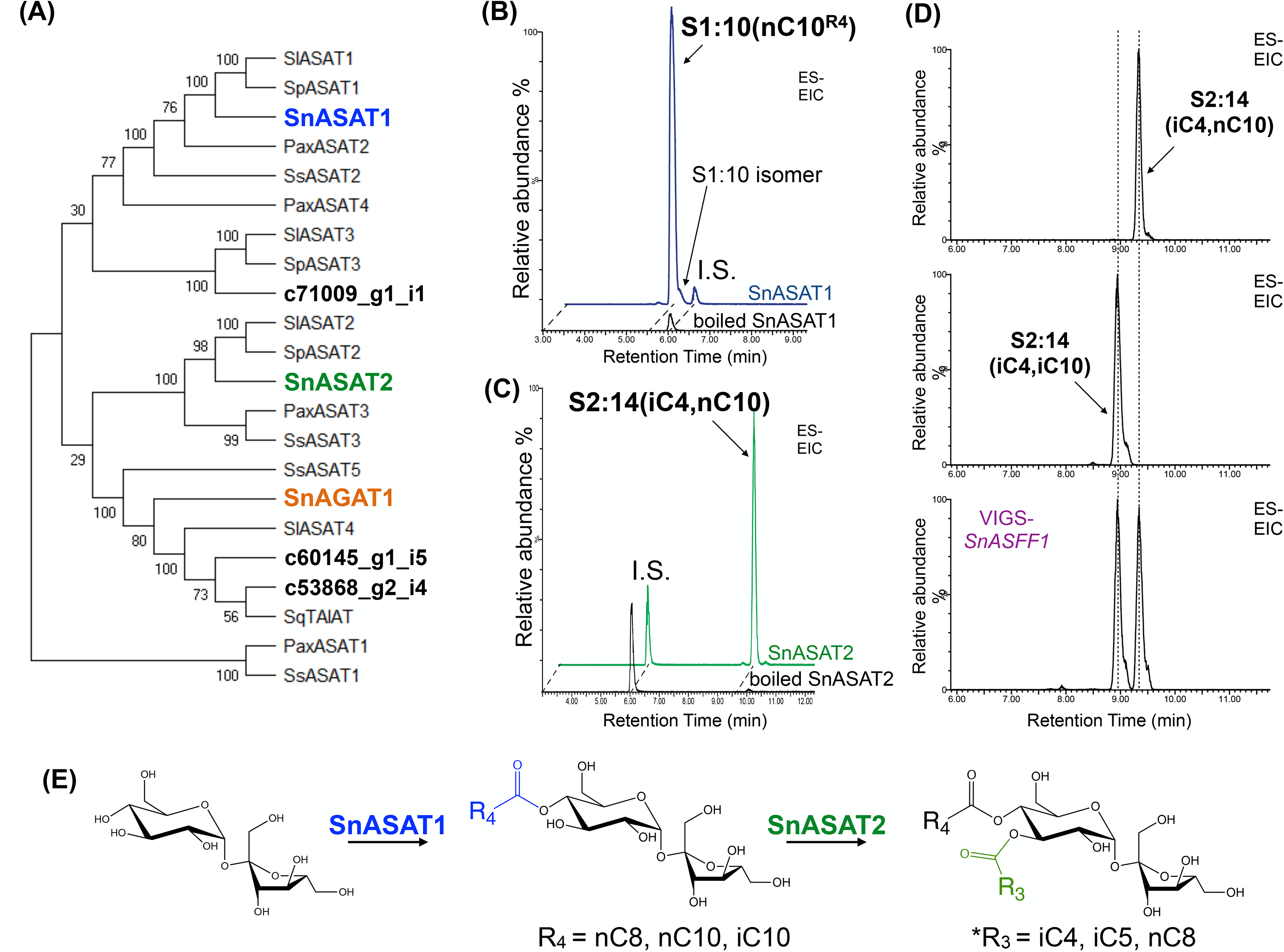
SnASAT1 and SnASAT2 sequential reaction produces diacylsucroses. (**A**) Maximum likelihood phylogeny analysis of amino acid sequences of ASAT candidates showed that SnASAT1 and SnASAT2 cluster with characterized ASAT1 and ASAT2 in tomato species, while SnAGAT1 clusters with the Solanaceae acylsugar acetyltransferases, SlASAT4, SsASAT5 and SqTAIAT. (**B**) LC-MS analysis of *in vitro* assay products showing that SnASAT1 produces monoacylsucrose S1:10(10^R4^) from sucrose and nC10-CoA substrates. The acylation at 4-position was verified by NMR as shown in Fig. S9. Blue trace represents full enzyme assays, whereas the black trace has heat-denatured SnASAT1. (**C**) LC-MS analysis of *in vitro* assay products showing that SnASAT2 produces diacylsucrose S2:14(4,10) from monoacylsucrose S1:10(10^R4^) and iC4-CoA substrates. Green trace represents full enzyme assays, whereas the black trace has heat-denatured SnASAT2. (**D**) *In vitro*-generated S2:14(iC4, nC10) and S2:14(iC4, iC10) co-elute (top and middle, respectively) with the *in vivo* S2:14(4,10) isomers from *SnASFF1*-silenced lines (bottom). S2:14(iC4, nC10) products in C and D are from two independent technical replicates. (**E**) Summary of the reactions catalyzed by SnASAT1 and SnASAT2 with sucrose and monoacylsucroses, respectively, and acyl-CoA substrates of different chain lengths. For more details, see Fig. S12. *Acylation at 3-position verified from SnASAT2-products co-elution with NMR characterized S2:14(4,10) with acyl chains as R_3_ and R_4_ from *S. nigrum*. All chromatographs are combined extracted ion chromatogram (ESI-) with telmisartan internal standard (I.S.) (*m/z* 513.23) and the formic adducts of S1:10(10) (*m/z* 541.25) or S2:12(4,10) (*m/z* 611.29).

We tested the hypothesis that these genes encode proteins of acylsugar biosynthesis by examining the activities of *Escherichia coli*-expressed and His-tag purified enzymes. C10, C8, iC5, iC4 and C2-CoAs were chosen as acyl chain donors in our *in vitro* assays, based on the abundance of these ester groups in our structural annotations (Fig. 1*B*). Using LC-MS, activity was detected with sucrose and the closest *S. nigrum* homolog of tomato ASAT1 (SlASAT1), which is encoded by the transcript *c63608_g1* (Fig. 4*B* and Fig. S8). This enzyme catalyzes the formation of monoacylsucroses – S1:8 and S1:10 – from sucrose and nC8, iC10 and nC10 acyl-CoAs (Fig. 4*B* and Fig. S8). The apparent *K_m_* values for SnASAT1 and nC8- and nC10-CoAs are 114 ± 21 and 11.7 ± 2.1 (µM), respectively (Fig. S9*A*). An apparent *K_m_* of 5 mM was measured for SnASAT1 and the acceptor substrate sucrose (Fig. S9*A*). Based on its *in vitro* activity and the *in planta* results presented below, we named this enzyme SnASAT1. SnASAT1-produced S1:10(nC10) co-eluted with SlASAT1-produced S1:10(nC10**^R4^**) but not with SsASAT1-produced S1:10(nC10**^R2^**) (Fig. S9*B*) (*10, 11, 13, 14*). Positive-mode MS fragmentation supports localization of the C10 chain on the six-member pyranose ring (Fig. S9*C*). This result was extended by NMR analysis, revealing S1:10(nC10^R4^) as the major product of SnASAT1 *in vitro* enzyme assays (Fig. 4*E* and Fig. S9*D*). A second, chromatographically separable S1:10 isomer, S1:10(10^R6^), accumulated in SnASAT1 enzyme assays (Fig. S10*A* and S9*D*). This 6-position acylated isomer appeared as a minor product in assays carried out at pH 6.0 with short incubation periods (5-30 min) (Fig. S10*A*). The concentration of this second isomer increases after extended incubation time, especially under neutral-to-alkaline pH conditions (Fig. S10*B*-*D*). For example, comparable amounts of S1:10(10^R4^) and S1:10(10^R6^) accumulated after 30 min at 30°C under pH 8.0 (Fig. S10*C*). We also observed non-enzymatic conversion of S1:10(10^R4^) to S1:10(10^R6^) after brief exposure to elevated temperature (65°C) (Fig. S10*B* and *D*) or extended incubation of purified S1:10(10^R4^) in unbuffered distilled water at room temperature. Non-enzymatic chain migration was previously documented for a R_4_ monoacylsucrose produced by SlASAT1, where the short acyl chain moved from R_4_ to R_6_ (*11*).

We characterized the activity of trichome-expressed *c65670_g1-*encoded SnASAT2, the closest homolog of tomato SlASAT2 and SpASAT2 (Fig. 4*A*). SnASAT2 utilized iC4-CoA as acyl-chain donor to decorate a second ring position on the pyranose ring of both S1:10(**n**C10^R4^) and S1:10(**i**C10) (Fig. 4*CD* and Fig. S11*A*). MS fragmentation patterns and retention times of the produced S2:14(iC4,iC10) and S2:14(iC4,nC10) (referred to below as ‘*in vitro* diacylsucroses’) were indistinguishable from the early- and late-eluting *in vivo* S2:14(4,10) metabolites, respectively (Fig. 4*D* and Fig. S11). We observed broad acyl donor and acceptor substrate specificity with iC4-, iC5- and nC8- serving as substrates with both S1:10(10^R4^) and S1:8(8) acceptors, producing diacylsucroses chromatographically indistinguishable from those detected in invertase-like deficient VIGS-*SnASFF1* lines (Fig. 4*E* and Fig. S12*A*). In contrast, no products of SnASAT2 were detected with unacylated sucrose, glucose, the SsASAT1 product S1:10(10^R2^), or the S1:10(10^R6^) SnASAT1 rearrangement product as acceptor (Fig. S12*B*-*D*). As seen with SnASAT1-generated monoacylsucroses, a second diacylsucrose isomer accumulated in a pH- and time-dependent manner after SnASAT1 + SnASAT2 reaction (Fig. S13). Together with the identification of SnASFF1, these results are consistent with the hypothesis that S2:14(4,10) from SnASAT1 + SnASAT2 sequential reactions are precursors of the *S. nigrum* di- and triacylglucoses (e.g. G2:14(4^R3^,10^R4^) and G3:16(2^R2^,4^R3^,10^R4^)).

### SnASFF1 is an acylsucrose hydrolase that evolved independently from SpASFF1

We tested SnASFF1 protein activity on *in vitro*- and *in vivo*-produced diacylsucroses, using recombinant His-tagged SnASFF1 protein expressed and purified from the *Nicotiana benthamiana* transient expression system(*15*). Indeed, SnASFF1 hydrolyzed all three tested diacylsucroses: the two *in vivo* pathway intermediates and the unrearranged *in vitro* diacylsucrose product generated by SnASAT1 and SnASAT2 sequential reactions (Fig. 5*A* and Fig. S14*A*). The reactions with *in vitro* and *in vivo* S2:14(4,10) isomers each yielded two major products that are the anomers of *S. nigrum* G2:14(4^R3^,10^R4^), as suggested by their identical retention times and mass spectra (Fig. 5*A* and Fig. S14*B*). In fact, the G2:14(4,10) isomers produced from *in vitro* S2:14(iC4,nC10) and *in vivo* late-eluting S2:14(4,10) are indistinguishable from the late-eluting anomers of *S. nigrum* G2:14(4^R3^,10^R4^). Additionally, SnASFF1 converted the *in vivo* early-eluting S2:14(4,10) into anomers co-eluting with those of the early-eluting *S. nigrum* G2:14(4^R3^,10^R4^) (Fig. 5*A* and Fig. S14*B*). We hypothesize that these two sets of diacylglucoses, differing in an nC10 or iC10 acyl chain as R_4_, collectively explain the overlapping signals of *S. nigrum* G2:14(4^R3^,10^R4^) (Fig. 5*A*). This result is consistent with the presence of abundant nC10 and iC10 in the acylsugar-derived ethyl ester profile (Fig. 1*B*). Taken together, these results provide strong evidence that SnASAT1, SnASAT2 and SnASFF1 are sufficient to reconstitute production of diacylglucoses produced in *S. nigrum* from sucrose and acyl-CoAs.

**Figure 5.**
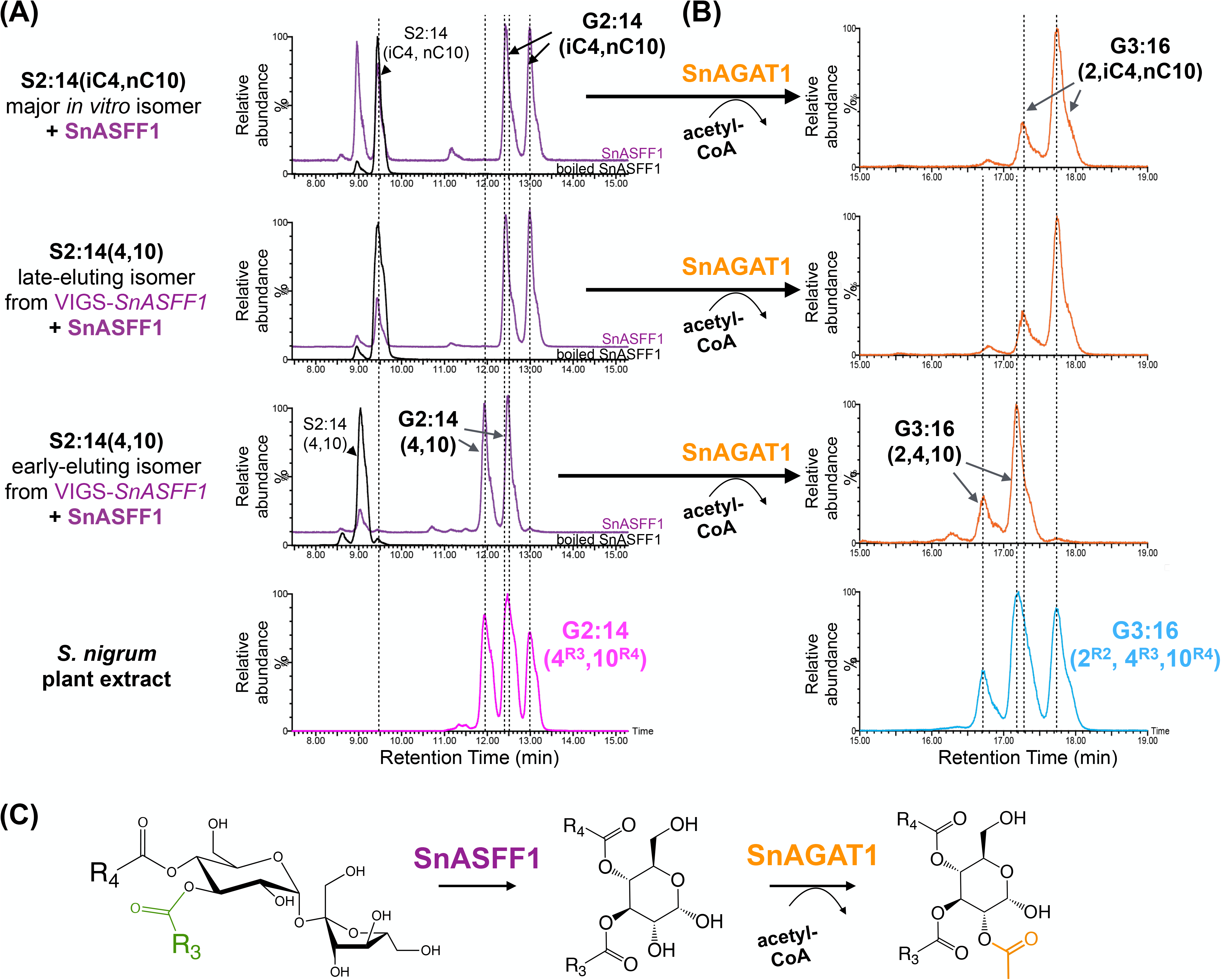
SnASFF1 and SnAGAT1 sequential activities with both *in vitro* and *in vivo* diacylsucrose substrates produces triacylglucoses. **(A)** SnASFF1 activity (purple traces) with the major *in vitro* S2:14(iC4, nC10) from consecutive SnASAT1 and SnASAT2 reactions (top); and the early- and late-eluting *in vivo* diacylsucroses S2:14(4,10) from VIGS-*SnASFF1* lines (bottom and middle, respectively) produced two distinct sets of diacylglucoses that collectively explains the multiple peaks of *S. nigrum* G2:14(4^R3^,10^R4^) (magenta trace). The diacylglucose products from *in vitro* S2:14(iC4, nC10) and the late-eluting *in vivo* S2:14(4,10) are chromatographically indistinguishable from the late-eluting *S. nigrum* G2:14(4^R3^,10^R4^). Black traces represent reactions with heat-denatured enzymes. **(B)** SnAGAT1 activities (orange traces) with the diacylglucoses produced by SnASFF1 activities with *in vitro-* and *in vivo* diacylsucrose S2:14(4,10) substrates produce two distinct set of triacylglucose products that collectively explains the triplet-like peaks of G3:14(2^R2^,4^R3^,10^R4^) in *S. nigrum* leaf extracts (cyan trace). Combined extracted ion chromatogram (EIC) under negative electrospray ionization (ESI-) showing formic adducts of S2:14(4,10) (*m/z* 611.29), G2:14(4,10) (*m/z* 449.24) and G3:16(2,4,10) (*m/z* 491.25).(**C**) Summary of results of SnASFF1 and SnAGAT1 activities in the reconstructed acylglucose biosynthetic pathway.

SnASFF1 hydrolytic activity was not detected using sucrose, in contrast to the canonical yeast invertase activity (Fig. S15*A*). Conversely, yeast invertase failed to use any tested S2:14(4,10) as substrate (Fig. S15*B*). These data are consistent with the hypothesis that SnASFF1 has acylsucrose-specific activity. We also observed SnASFF1 hydrolytic activity with *in vitro*-produced S1:10(10^R4^) and the *S. pennellii* triacylsucrose S3:18(4^R2^,4^R4^,10^R3^) as substrate (Fig. S15*B*). However, the monoacylglucose product, G1:10(10), does not appear to be an acceptor substrate for SnASAT2 with nC10-, nC8-, iC5-, iC4- and acetyl-CoA (Fig. S12*A*). Taken together, the results support the hypothesis that SnASFF1 is an acylsucrose hydrolase that contributes to *S. nigrum* acylglucose biosynthesis by converting diacylsucroses to diacylated glucoses.

To explore the evolution of acylglucoses in *S. nigrum* and wild tomato, we generated a gene tree with invertase-like genes and transcripts from *S. lycopersicum*, *S. pennellii* and *S. nigrum* (Fig. 6). Phylogenetic analysis showed that the *S. nigrum ASFF1* is a close homolog of the tomato *LIN5* (Solyc09g010080) and *LIN7* (Solyc09g010090) paralogs – tandem duplicates of sucrose-specific invertases. In contrast, the *S. pennellii ASFF1* is in a clade distinct from *SnASFF1*, with greater similarity to tomato Solyc03g121680 and Solyc06g064620 and the two *S. nigrum* transcripts, *c65240_g1* and *c62944_g1*. We observed no acylsugar phenotype upon *in planta* silencing *c65240_g1,* the closest BLAST hit of SpASFF1 in *S. nigrum* transcriptome (Fig. S15*B*). This result is consistent with the low *S. nigrum* trichome expression of *c65240_g1*. Together, these relationships indicate that the acylsugar hydrolyzing activity of SnASFF1 and SpASFF1 evolved independently in *S. nigrum* and the wild tomato, *S. pennellii*.

**Figure 6.**
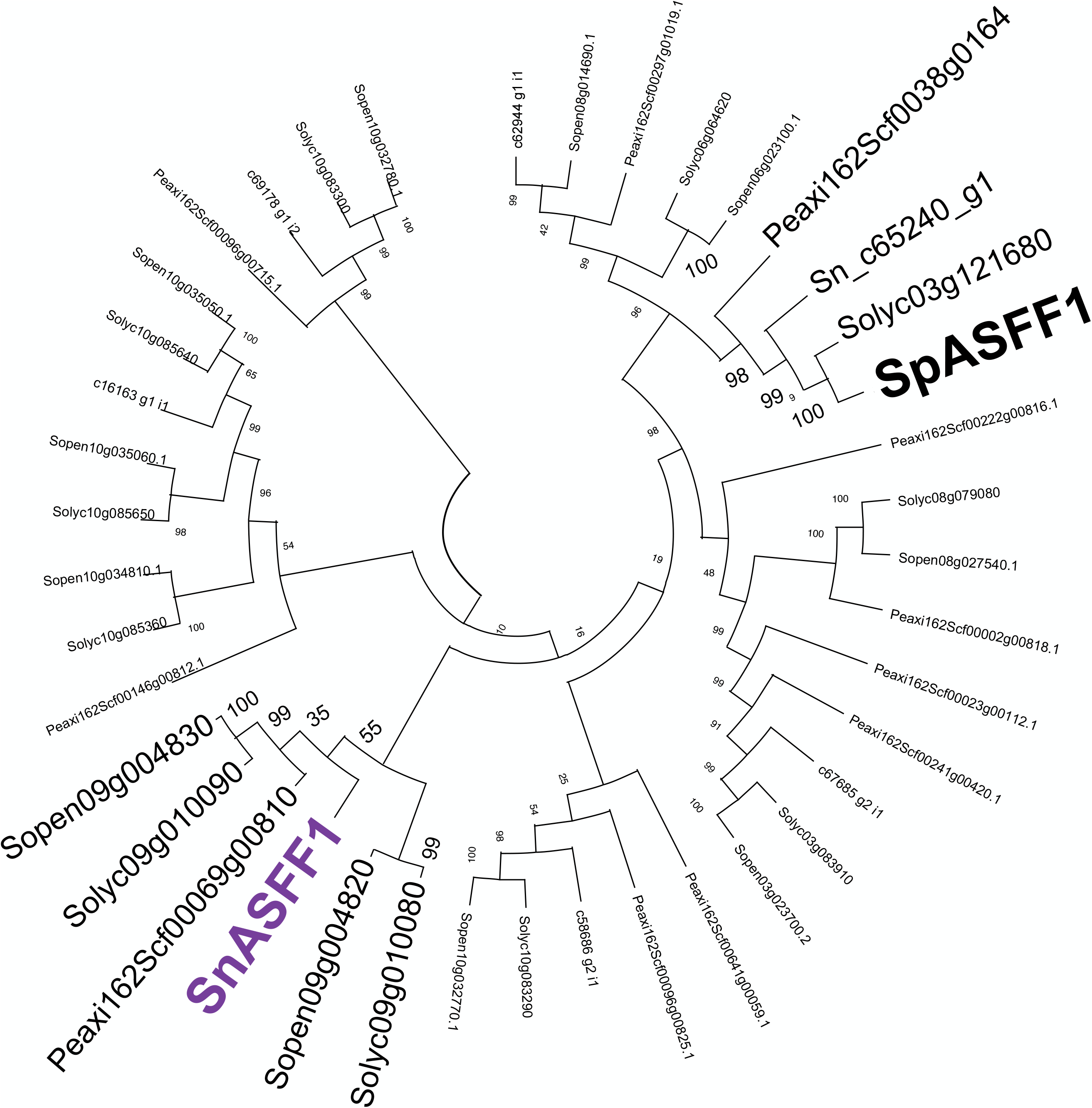
Phylogenetic analysis of SnASFF1 and SpASFF1 reveals independent evolution. Invertase-like proteins from *S. lycopersicum*, *S. pennellii* and *S. nigrum* were aligned using MUSCLE with default parameters in MEGA X. SnASFF1 is marked bold in purple, whereas SpASFF1 is marked bold in black. Maximum likelihood tree constructed using the Jones-Taylor-Thornton algorithm with 1000 bootstrap support using the topology-only tree in MEGA X.

### Identification of a trichome-specific BAHD acylglucose acetyltransferase

The *in vitro* production of diacylglucoses by sequential reaction of SnASAT1, SnASAT2 and SnASFF1 is consistent with the hypothesis that the acetylation step leading to *S. nigrum* triacylglucoses is carried out on diacylglucose substrates. Indeed, the search among the remaining trichome-enriched ASAT candidates led to identification of acetylation by SnAGAT1 (***S****. **n**igrum* **a**cyl**g**lucose **a**cyltransferase 1; encoded by *c65306_01*), as the last step of triacylglucose biosynthesis in *S. nigrum*.

As hypothesized, SnAGAT1 acetylated the diacylglucoses produced by SnASFF1 activities with *in vitro* and *in vivo* diacylsucrose S2:14(4,10) substrates (Fig. 5*B*). The G3:16(2,4,10) anomers obtained from both *in vitro* S2:14(iC4,nC10) and *in vivo* late-eluting S2:14(4,10) have retention times and mass spectra identical to the late-eluting peaks of *S. nigrum* G3:16(2^R2^,4^R3^,10^R4^) (Fig. 5*B* and Fig. S16*A*). SnASFF1 and SnAGAT1 sequential activities with *in vivo* early-eluting S2:14(4, 10) – which shares identical retention times and mass spectra with *in vitro* generated S2:14(iC4,iC10) – generated early-eluting triacylglucose anomers (Fig. 5*B* and Fig. S16*A*). We hypothesize that these two sets of triacylglucoses, with an nC10 or iC10 chain, collectively explain the multiple peak signals of *S. nigrum* G3:16(2^R2^,4^R3^,10^R4^) (Fig. 5*B* and Fig. S3). This result is consistent with the abundance of nC10 and iC10 acyl chains on *S. nigrum* acylsugars (Fig. 1*B*). We also observed SnAGAT1 activity with the diacylglucose-rich *S. nigrum* extract fraction 2 as acceptor substrates and acetyl-CoA as donor, leading to accumulation of triacylglucoses that are indistinguishable to *S. nigrum* triacylglucoses (Fig. S16*C*). These assays provide strong evidence that SnAGAT1 catalyzes the last step in *S. nigrum* triacylglucose production (Fig. 5*C*).

### *In vitro* validation of the *S. nigrum* acylglucose biosynthetic pathway

We independently assessed the functions of the three BAHD enzymes on the reconstructed pathway by taking advantage of BAHD acyltransferase reversibility in the presence of free coenzyme A. Using plant-derived acylglucoses as substrates for reverse enzyme assays, we verified that SnAGAT1 and free coenzyme A converts *S. nigrum* extract fraction 1 triacylglucoses into diacylglucoses (Fig. 7*A*). The ion masses and mass spectra of these diacylglucoses are consistent with loss of an acetyl group (Fig. S16*B*), supporting the designation of SnAGAT1 as an acetyltransferase (Fig. 5*C* and 7*B*).

**Figure 7.**
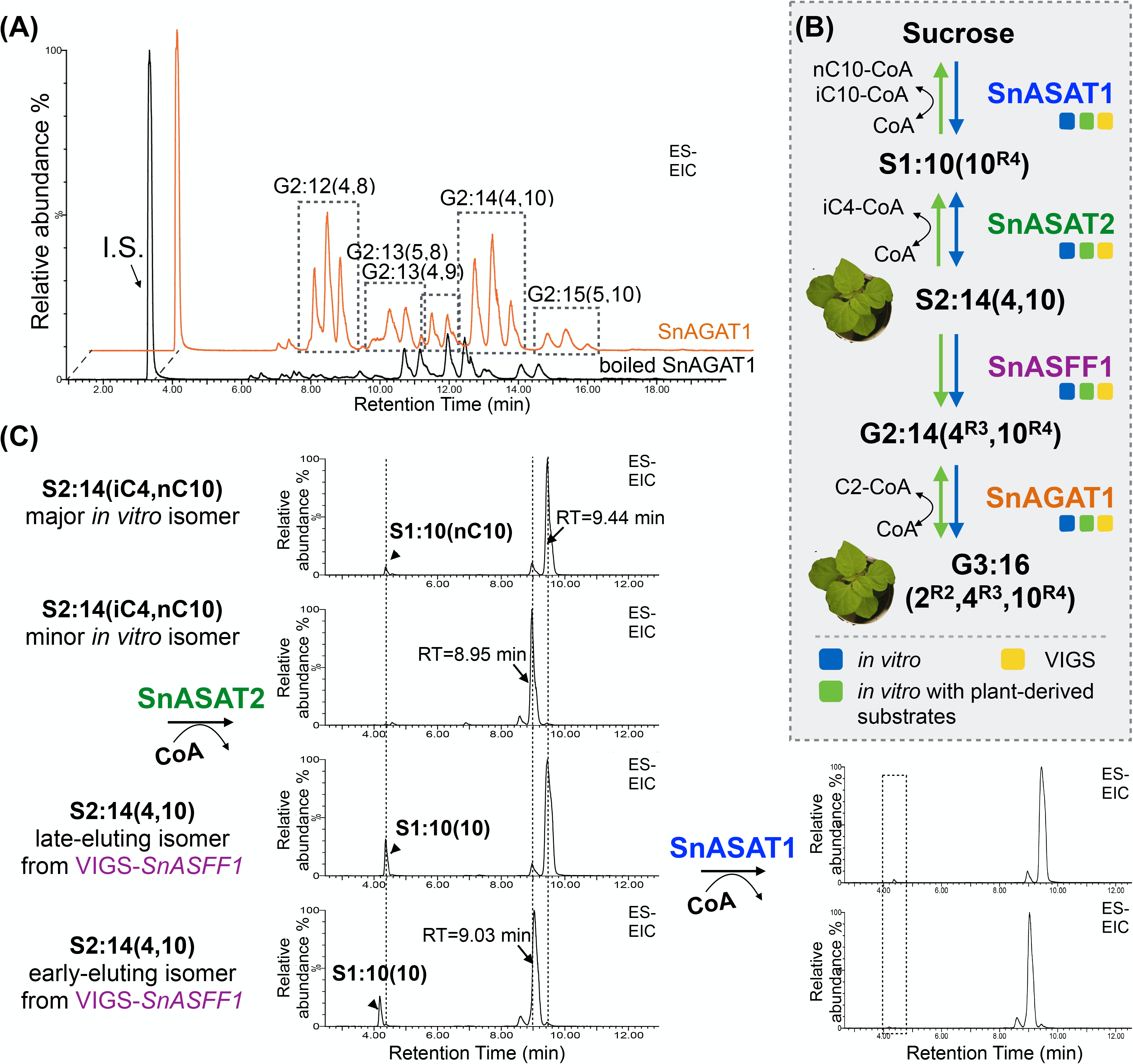
Independent validation of the reconstructed acylglucose biosynthetic pathway with *in vitro* reverse activities. **(A)** LC-MS analysis of reverse enzyme assay products from SnAGAT1 activity (orange trace) shows SnAGAT1 deacetylating *S. nigrum* fraction 1 tri-acylglucoses in the presence of free coenzyme A. Combined extracted ion chromatogram (EIC) under negative electrospray ionization (ESI-) showing telmisartan as internal standard (I.S.) (*m/z* 513.23) and formic adducts of G2:12, G2:13, G2:14 and G2:15. The corresponding *m/z’s* of these compounds are listed in Table S1. **(B)** Schematic representation of the reconstructed *S. nigrum* triacylglucose biosynthetic pathway. Blue arrows indicate *in vitro* pathway reconstruction starting from sucrose, green arrows indicate enzyme activities with plant-derived products. **(C)** SnASAT2 activities with the major (top) – but not minor (second from the top) – *in vitro* S2:14(iC4, nC10) products and the early- and late-eluting *in vivo* diacylsucrose S2:14(4,10) substrates from VIGS-*SnASFF1* lines (second from the bottom and bottom, respectively) produce monoacylsucrose products that can further be deacylated by SnASAT1 activity (bottom right). The monoacylsucrose intermediates from the late-eluting *in vivo* S2:14(4,10) and *in vitro* S2:14(iC4, nC10) (top and third from top) are chromatographically indistinguishable from S1:10(nC10^R4^). Combined extracted ion chromatogram (EIC) under negative electrospray ionization (ESI-) showing formic adducts of S2:14(4,10) (*m/z* 611.29) and S1:10(10) (*m/z* 541.25).

SnASAT2 was tested for reverse activity with both early- and late-eluting *in vivo* S2:14(4,10) from VIGS-*SnASFF1* lines and the major and minor SnASAT1 + SnASAT2 *in vitro*- generated S2:14(iC4,nC10) isomers (Fig. 7*C*). SnASAT2 converted the *in vivo*-derived and the major *in vitro*-produced diacylsucroses to monoacylsucroses S1:10(10) (Fig. 7*C*). The monoacylsucroses produced from both *in vitro* S2:14(iC4,nC10) and *in vivo* late-eluting S2:14(4,10) are chromatographically indistinguishable from the SnASAT1-forward reaction product S1:10(nC10^R4^) (Fig. 7*C*). Addition of SnASAT1 to either of the SnASAT2 reverse reaction assays reduced the accumulation of S1:10(10) (Fig. 7*C*). In contrast, no reverse activity was detected using the minor *in vitro* diacylsucrose rearrangement product as substrate (Fig. 7*C*). Similarly, reverse assays using *in vitro*-generated monoacylsucroses as substrates revealed SnASAT1 enzymatic activity with S1:10(10^R4^), but not with the S1:10(10^R6^) rearrangement product (Fig. S17). Together, these data provide independent evidence for the *S. nigrum* acylglucose biosynthetic pathway, with SnASAT1, SnASAT2, SnASFF1 producing diacylglucoses, and SnAGAT1 catalyzing the final step in triacylglucose biosynthesis (Fig. 7*B*).

### VIGS validation of three acyltransferases in *S. nigrum* acylglucose biosynthesis

We deployed VIGS to test the hypothesis that SnASAT1, SnASAT2 and SnAGAT1 activities are necessary for *in vivo* acylglucose biosynthesis (Fig. 8). As expected for early steps in the core acylsugar biosynthetic pathway, silencing *SnASAT1* and *SnASAT2* each led to reduction of the four most abundant di- and triacylglucoses (Fig. 8*A* and *B*, Fig. S18*A* and *B* and Table S6 and S7). Silencing SnAGAT1 caused statistically significant increases in the ratio of diacylglucoses to corresponding triacylglucoses relative to the controls (Fig. 8*C*, Fig. S18*C* and Table S8). These results provide strong support that di- and triacylglucoses are produced *in vivo* via the three- and four-step biosynthetic pathway demonstrated through *in vitro* biochemistry, respectively (Fig. 7*B*).

**Figure 8.**
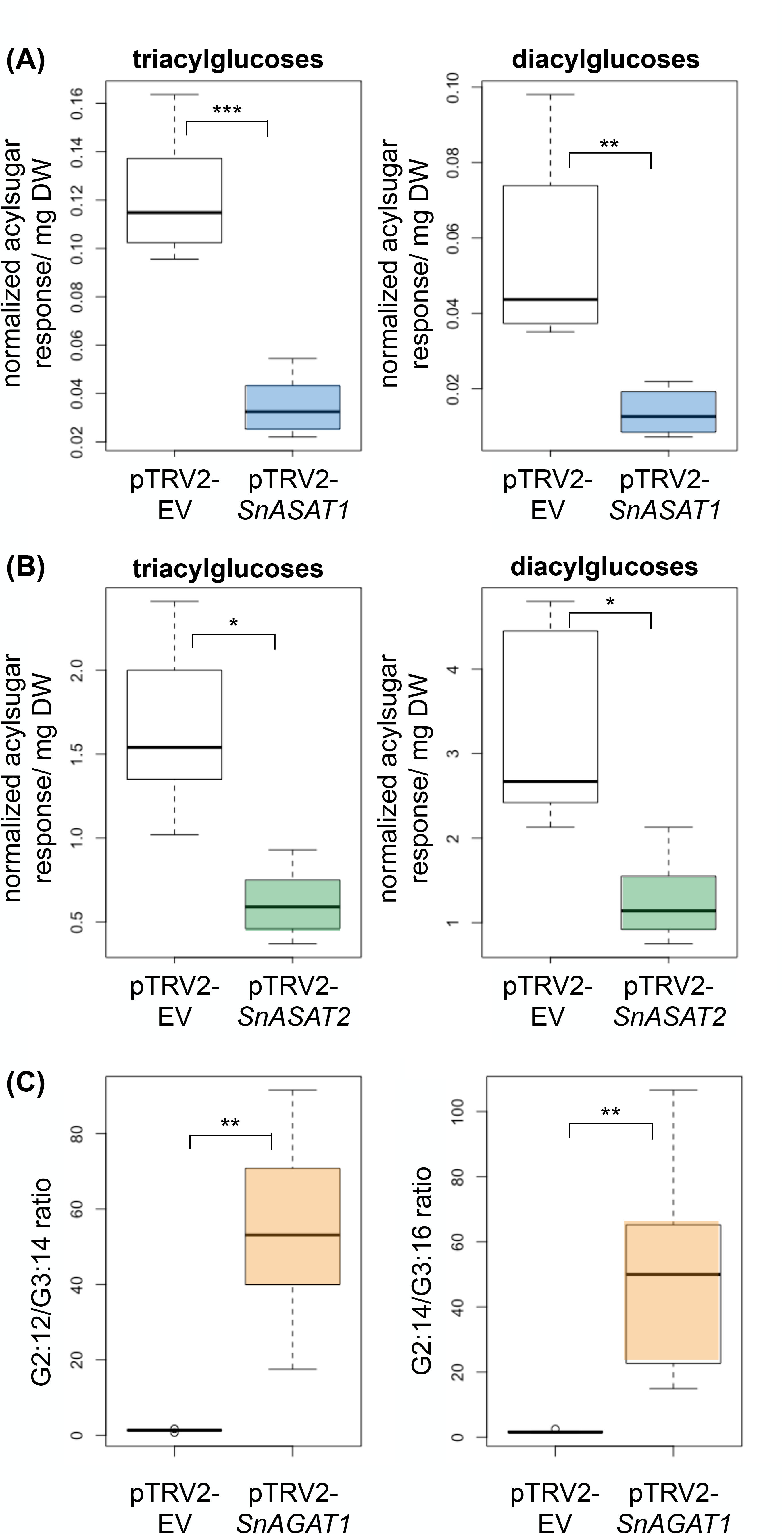
*In vivo* validation of the reconstructed acylglucose biosynthetic pathway with VIGS. **(A)** Comparison of acylglucose accumulation in *SnASAT1*-targeted and empty vector VIGS plants. Acylsugars were analyzed using LC-MS in ESI+ mode. (**B**) Comparison of acylglucose accumulation in *SnASAT2*-targeted and empty vector VIGS plants. Acylsugars were analyzed using LC-MS in ESI- mode. **(C)** Comparison of the ratio of diacylglucose G2:14 and triacylglucose G3:16 in *SnAGAT1*-targeted and empty vector VIGS plants. Diacylglucose quantities were measured by integrating peak areas of G2:12, G2:13, G2:14 and G2:15, whereas triacylglucose quantities were measured by integrating peak areas of G3:14, G3:15, G3:16 and G3:17. The integrated peak areas were normalized to the internal standard telmisartan and dry leaf weights. Peak areas of G2:14 and G3:16 were integrated under negative mode and normalized to the internal standard telmisartan and dry leaf weights to measure the ratio between tri- and diacylglucoses. The corresponding *m/z* of analyzed acylsugars are listed in Table S1 unless otherwise specified. Significant levels are shown (*, *p* < 0.05; **, *p* < 0.01; ***, *p* < 0.001; Welch’s two sample t-test).

## Discussion

While evolutionary diversification occurs by relatively simple mechanisms of gene duplication, changes in gene expression, and modification of gene product activities, the resulting phenotypic diversity is remarkable. Plant specialized metabolism is a poster child for how a matrix of a relatively small number of chemical feedstocks – generally from core metabolism – and varied enzyme classes, lead to hundreds of thousands of metabolites of diverse structure and function. The sheer number and structural variety of these compounds presents challenges for the structural elucidation essential for rigorous metabolic pathway dissection. Conversely, the ability to use *in vitro* pathway reconstruction to test hypotheses regarding pathway evolution is a strength of evo-metabolism.

Acylsugar metabolism has become an exemplary system for understanding both the general principals and specific mechanisms by which enzymes are repurposed and combined to generate phenotypic diversity. Here we describe three characteristics of *S. nigrum* acylsugar diversity and metabolism not previously reported for other species. First, although only acylhexoses were detected, they include a mixture of both inositol and glucose esters (Fig. 1 and 2). Second, an ***acylglucose*** acyltransferase is involved in making triacylglucoses (Fig. 5). Finally, the acylglucose pathway intermediates are produced from a neofunctionalized invertase, which evolved independently from the recently reported wild tomato SpASFF1 triacylsucrose β-fructofurosidase (Fig. 5 and 6). This work was facilitated by integration of LC- and GC-MS combined with 2-D NMR methods, bypassing a requirement for metabolite purification (Fig. 1).

### Integrative approaches to elucidate the *S. nigrum* acylglucose biosynthetic pathway

We present four lines of evidence that *S. nigrum* di- and triacylglucoses are synthesized by SnASFF1 and SnAGAT1 from the diacylsucrose intermediates produced by sequential reactions of SnASAT1 and SnASAT2. First, we reconstructed the biosynthetic pathway *in vitro* using purified enzymes, sucrose and acyl-CoAs (Fig. 4 and 5). The acylglucoses generated are chromatographically indistinguishable from those observed in *S. nigrum*, while intermediates of the reconstructed pathway have acylation positions consistent with the *in vivo* final products. Second, genetic evidence supports the involvement of these enzymes in *S. nigrum* acylglucose biosynthesis (Fig. 3 and 8): VIGS silencing of the BAHD acyltransferases *SnASAT1* and *SnASAT2* reduces total acylglucoses*, SnAGAT1* reduces total triacylglucoses, while *SnASFF1* silencing causes accumulation of diacylsucrose pathway intermediates. Third, we cross-validated that the major *in vitro* diacylsucrose intermediates produced by the reconstructed pathway behave chromatographically and biochemically identically to *in vivo* diacylsucrose intermediates purified from VIGS-*SnASFF1* lines (Fig. 4*D*, 5 and 7C). Notably, SnASFF1 converted both *in vivo* and *in vitro* synthesized diacylsucroses to diacylglucose products that can be converted to triacylglucoses by SnAGAT1. In addition, both the di- and triacylglucose anomers produced are indistinguishable from those extracted from *S. nigrum*. While this biochemical and genetic evidence provided strong support for the reconstructed pathway, the observation of mono- and diacylated *in vitro* rearrangement products under pH >6 and elevated temperature (Fig. S10 and S13) led us to seek a fourth line of evidence for pathway validation using reverse BAHD assays.

We compared reverse enzymatic products of *in vitro* and *in vivo* intermediates to verify the relevance of all the *in vitro*-produced intermediates. SnASAT2-reverse activity deacylates the *in vivo* – and major *in vitro* – diacylsucroses to monoacylsucroses that are indistinguishable from those obtained from SnASAT1 forward assays. All of these monoacylsucrose products are substrates for SnASAT1 reverse activities, supporting the hypothesis that SnASAT1 acylates at the 4-position. The lack of SnASAT2 reverse enzyme activity with *in vitro* minor diacylsucrose is consistent with the hypothesis that the rearranged product is not a biosynthetic intermediate. This result is consistent with NMR analysis suggesting that the 6-position acylation of the medium-length chain on the minor monoacylsucrose products is not observed on *S. nigrum* acylglucoses. Hence, we conclude that the minor *in vitro* artifacts are not intermediates of the *S. nigrum* acylglucose biosynthetic pathway, despite their similar – but not identical – retention times and mass spectra.

The reverse activity also enabled us to facilitate pathway discovery. Using abundant plant-derived products, we discovered the first reported acylglucose acyltransferase, SnAGAT1, acting as the last step in *S. nigrum* triacylglucose biosynthesis (Fig. 7*A*). This acetyltransferase activity was later verified by forward enzyme assays with acetyl-CoA and *in vitro* diacylglucoses produced by the multistep – SnASAT1 + SnASAT2 + SnASFF1 – reactions (Fig. 5*B*). SnAGAT1 is the third phylogenetically related acetyltransferase in acylsugar biosynthesis, falling in the same BAHD cluster with tomato SlASAT4, *S. quitoense* TAIAT and *S. sinuata* SsASAT5 (Fig 3*A*). Deploying BAHD reverse enzyme assays accelerated SnAGAT1 identification and allowed *in vitro* pathway reconstruction in the non-model species. In fact, the *S. nigrum* acylglucose biosynthetic pathway is the first completely *in vitro* reconstructed acylhexose pathway in a non-model organism without the powerful genetic resources that enabled the discovery of *S. pennellii* acylglucose biosynthetic pathway.

### Convergent evolution of acylsucrose hydrolyzing enzymes

Phylogenetic analysis of SnASFF1 and restricted accumulation of acylhexoses across the Solanaceae is consistent with the hypothesis that acylglucose production arose independently in the Old World black nightshade *S. nigrum* and New World wild tomato *S. pennellii* (Fig. 6). One strong line of evidence is that the two ASFF enzymes are members of different clades within the β-fructofuranosidase gene tree (Fig. 6). Second, SpASFF1 – and synthesis of its P-type triacylsucrose substrates (*15*) – are restricted within a subclade of the tomato Solanum group (*12*), consistent with this being a derived trait within the New World Solanum. The independent evolution of acylhexose biosynthesis raises the intriguing question of whether there are phenotypic advantages of producing acylsugars with a glucose core. Similarly, the observation of mixed acylsucroses and acylglucoses in *S. pennellii*, and mixed acylglucose and acylinositols in *S. nigrum* begs the question of whether there are synergistic effects of acylglucoses with other acylsugars.

Despite co-option from distinct lineages, the two independently evolved ASFFs are both trichome-enriched, neofunctionalized GH32 β-fructofuranosidases, which lack the canonical **D**DT**K** sucrose-binding pocket of Arabidopsis Cell-Wall Invertase 1 (ACWINV1; AT3G13790) (*37–39*) (Fig. S4). This is consistent with the observation that neither of the ASFFs utilize un-acylated sucrose as substrate (*15*) (Fig. S12*C*). In addition, both ASFFs are expressed in the same tip cells as their acylsucrose substrates (*15, 40*). It will be interesting to learn whether invertase-like enzymes evolved to produce the acylglucoses reported in *Datura* and *Nicotiana* species (*21, 22*). The diversity of apparent *in vivo* substrates for both ASFFs – diacylsucroses for SnASFF1 and triacylsucroses for SpASFF1 – also present opportunities for enzyme structure and function analysis.

### Streamlined analytical approaches for metabolite annotation

It is remarkable that we continue to discover metabolic variations as new Solanaceae species are analyzed. Because biosynthetic pathway dissection requires knowledge of product and intermediate structures, this process can be a bottleneck. We developed a structural analysis pipeline that reduces the cost and time involved in sample purification. The presence of three distinct acylhexose classes – di- and triacylglucoses and triacylinositols – in *S. nigrum* presented a challenging case study for this integrative approach. We designed the pipeline to leverage the substantial chemical shift library assembled from acylsugar studies in other Solanaceous plants (*10, 11, 13, 29, 34, 41*).

While the pipeline performed as expected for the most abundant triacylsugars in *S. nigrum*, the many overlapping NMR signals in total trichome acylsugar extracts led us to perform partial purification using silica gel chromatography. We obtained clear signals of *S. nigrum* triacylglucoses from the less polar fraction, while 2D HSQC-TOCSY successfully separated proton correlation signals from diacylglucoses and triacylinositols in the high polarity fraction. With the simple partial purification, our NMR approach provided sufficient information to address key characteristics of the enzymes involved in acylsugar biosynthesis without the need to purify individual compounds: (1) recognition of different acylated sugar cores in mixtures and (2) assignments of ester linkage positions on sugar cores. This approach extends the use of 2D ^1^H-^13^C HSQC NMR spectra beyond previously published characterization of lignin structures (*42*). Limiting the need for purification is particularly beneficial for investigating anomers and isomers differing in acyl chain branching patterns, where overlapping retention times can frustrate attempts at large-scale purification to homogeneity. As acylsugars are sporadically found in Martyniaceae, Rosaceae, Geraniaceae, Caryophyllaceae and Brassicaceae plants, our approach can further assist with large scale screening for sugar esters beyond Solanaceae (*43–47*). The vast complexity of specialized metabolism in the plant kingdom will be more efficiently addressed by exploiting the variety of multidimensional NMR strategies to aid in metabolite annotations from spectra of mixtures.

In addition to illustrating principles by which the evolution of form occurs, our deepening understanding of specialized metabolic innovation has practical benefits. Metabolic engineering continues to be a trial and error enterprise (*48*). Understanding the successful evolutionary outcomes leading to synthesis of biologically active metabolites provides new enzymes, transporters and transcriptional regulators for the synthetic biology toolkit. Documenting recurring themes, such as the independent recruitment of invertases in multiple acylglucose-producing *Solanum* lineages, indicates robust strategies for engineering synthesis of these protective compounds and other structurally similar compounds of economic or pharmaceutical importance.

## Materials and Methods

### Plant Material

Seeds of *S. nigrum* were obtained from New York Botanical Garden(*14*). For germination, seeds were treated with half-strength bleach for 5 min and rinsed six times in deionized water before sowing on moist filter paper in petri dishes at 28°C. Seedlings were transferred to Jiffy-7® peat pellets (Jiffy Products of America, OH, USA) upon germination. Plants used for analysis were grown in a growth chamber at 22°C under a 16-hour photoperiod (70 µmol m^−2^ s^−1^ photosynthetic photon flux density) with relative humidity set to 50%.

### Acylsugar metabolite annotation

The acylsugar extraction protocol for LC-MS analysis is available in Protocols.io at https://dx.doi.org/10.17504/protocols.io.xj2fkqe. As described previously in Leong et al. (2019), leaf surface acylsugar extraction were carried out by gently agitating a single leaflet in 1 mL acetonitrile:isopropanol:water (3:3:2 v/v with 0.1% formic acid and 1 µM telmisartan; isopropanol obtained from J.T.Baker, Phillipsburg, NJ, USA, all others obtained from Sigma-Aldrich, St. Louis, MO, USA) for 2 min. Telmisartan acts as an internal standard for high-performance liquid chromatography (HPLC). The extraction solvent was collected and stored in 2-mL LC-MS vials at -20°C. All extracts were analyzed on LC-MS (Waters Corporation, MA, USA) using 7-min, 30-min or 110-min LC gradients on an Ascentis Express C18 HPLC column (10 cm x 2.1 mm, 2.7 µm) (Sigma-Aldrich, St. Louis, MO, USA), which was maintained at 40°C. The 110-min method minimized chromatographic overlap in support of metabolite annotation. The HPLC-MS methods are described in protocols.io and Table S10.

Acylsugars structures were inferred by positive and negative mode MS collision-induced dissociation as described previously (*11, 29, 32, 41*). In short, co-eluting fragments generated by collision-induced dissociation in negative ion mode were compared among three energy potentials to confirm acylsugar metabolites. Annotation strategy, thresholds and confidence levels were described in detail in Supplementary Text.

Acylsugar acyl chain composition was determined by ethyl ester derivatization and subsequent GC-MS analysis. To create fatty acid ethyl esters, acylsugar samples were saponified and transesterified as previously described in Ning et al. (2015) with some modification. In short, a leaflet was immersed in 1 mL of acetonitrile/isopropanol (1:1 v/v) for 2 min with gentle agitation. The extracts were transferred, evaporated to dryness under flowing air and redissolved in ethanol with 300 µL of 21% (v/v) sodium ethoxide (Sigma-Aldrich, St. Louis, MO, USA). The reaction was gently vortexed every 5 min for 30 min in the fume hood. Four hundred µL hexane with 55 µg/mL of tetradecane (internal standard; Sigma-Aldrich, St. Louis, MO, USA) was added to the reaction mixture for phase separation. The hexane layer was transferred and extracted three times using 500 µL saturated aqueous sodium chloride each time. The final hexane phase (∼100 µL) was transferred to autosampler vials with glass inserts and analyzed by capillary GC-MS on an Agilent J&W DB-5 column (10-m, 0.1-mm (i.d.) fused silica column with a 0.34-µm-thick stationary phase; Agilent). One microliter of each hexane extract was injected using splitless mode. The gas chromatography program is described in Table S10. All compounds were analyzed using an Agilent 6890N gas chromatograph/Agilent 5975B single quadrupole mass spectrometer using 70 eV electron ionization. Fatty acid ethyl esters were identified by library search against Agilent RTL library and compared with commercially available standards.

For total extract NMR profiling, acylsugars were extracted from fifteen 3-4 weeks old *S. nigrum* plants by dipping aerial tissue into 500 mL of ethanol with 0.1% (v/v) formic acid with gentle agitation in a 1-L beaker. The ethanol was evaporated under reduced pressure using a rotary evaporator with a warm water bath (40°C). The dried residue (∼5 mg) of *S. nigrum* surface plant extracts were dissolved in CDCl3 (99.8 atom % D-, Sigma Aldrich, St. Louis, MO, USA) and transferred to solvent-matched (5 mm) NMR Shigemi tube (Shigemi Co., LTD., Tokyo, Japan) for analysis. ^1^H, HSQC, HMBC, and TOCSY spectra were recorded using the Avance 900 MHz spectrometer (Bruker, Billerica, MA, USA) equipped with a TCI triple resonance probe at the Michigan State University Max T. Rogers NMR facility (See Table S2-4 for more details). All spectra were referenced to non-deuterated CDCl3 solvent signals (δH = 7.26 (s) and δC = 77.2 (t) ppm).

For silica gel column chromatography, *S. nigrum* surface plant ethanol extract was concentrated *in vacuo* (∼8 mg), dissolved in ethyl acetate/ hexane (1 mL, 1:1 v/v) and loaded to the silica gel column (100 g, 200-425 mesh, 60 A, Jade Scientific Inc, Westland, MI, USA) packed with ethyl acetate/hexane (200 mL, 1:1 v/v) slurry. The compounds were eluted using mobile phase ethyl acetate/hexane (400 mL, 2:1 v/v in 0.02% acetic acid) with compressed air flash column chromatography. After TLC analysis (4:1 ethyl acetate/hexane, p-Anisaldehyde stain), fractions *F*15-17 and *F*24–28 were combined, concentrated *in vacuo* to give fraction 1 (∼2 mg, Rf = 0.7) and fraction 2 (∼2 mg, Rf = 0.3). Combined samples were analyzed on NMR as described above.

### Gene Identification and Phylogenetic Analysis

All transcript assemblies and expression data are from Moghe et al. (2017) and were analyzed using Geneious R9.1.8 and R, respectively. To identify BAHD candidates, BLAST and TBLASTN searches were performed using ASAT sequences from tomato and *S. sinuata*. The TBLASTN hits were parsed to include only those with HXXXD motifs and DFGWG-like motif (one mismatch). The trichome-stem expression data of the remaining sequences were obtained from expression data available in the study by Moghe et al. (2017), whereas a length of 400 to 500 amino acids and the relative positions of the two motifs on these sequences were confirmed manually(*36*). To identify ASFF candidates, 23 invertase-like sequences from tomato were used to BLAST and TBLASTN search *S. nigrum* transcriptome. The obtained sequences were checked for trichome-enrichment and WXNDPNG, RDP, EC and DXXK motifs as described above.

Phylogenetic reconstructions were performed using MEGA X(*49*). To obtain the BAHD tree, BAHD candidate sequences were aligned against several characterized ASATs(*9–11, 14*) and several other BAHD sequences from D’Auria (2006) using the MUSCLE algorithm under default parameters. To obtain the invertase tree, invertase-like proteins from *S. lycopersicum*, *S. pennellii* and *S. nigrum* were aligned with the same algorithm and settings. Maximum likelihood estimations were performed with Jones-Taylor-Thornton (JTT)+G+I with 5 rate categories as the substitution model. One thousand bootstrap replicates were performed using partial deletion (30% gaps) for tree reconstruction.

### Transient expression and purification of BAHD protein

All Sanger DNA sequencing confirmations in this study were performed with the indicated sequencing primers at the Research Technology Support Facility Genomics Core, Michigan State University, East Lansing, MI. All primer sequences are listed in Table S9.

Recombinant BAHD proteins were generated using *E. coli* as the host for enzyme assays. The full-length open reading frames of *c63608_g1*, *c65670_g1*, *c71009_g1*, *c53868_g2*, *c60145_g1*, *c65306_g1* were amplified from *S. nigrum* young leaf and peduncle cDNA and cloned into pET28b(+) (EMD Millipore, MA, USA) using BamHI and XhoI restriction sites and 2× Gibson Assembly master mix (NEB, Ipswich, MA, USA) according to the manufacturer’s instructions. The assembled constructs were transformed into BL21 Rosetta (DE3) cells (EMD Millipore, MA, USA) and submitted for Sanger sequencing.

Protein expression was carried out as described before (*11, 16*). In short, 1 L Luria-Bertani (LB) media with kanamycin (50 µg mL^-1^) and chloramphenicol (33 µg mL^-1^) were inoculated 500:1 with an overnight culture obtained from the bacterial strain with the desired construct. After incubating at 37°C, 225 rpm and reaching OD600 between 0.5 and 0.6, large scale cultures were chilled on ice for 20 min before a final concentration of 50 µM isopropylthio-β-galactoside was added. Cultures were incubated at 16°C and 180 rpm for 16 h before cell pellets were harvested by centrifuging at 4,000g for 10 min under 4°C. The following protein extraction steps were also performed on ice or at 4°C. The cell pellets were resuspended in 25 mL of extraction buffer (50 mM NaPO_4_, 300 mM NaCl, 20 mM imidazole, and 5 mM 2-mercaptoethanol, pH 8) by vortexing and submitted to eight cycles of sonication (30 s on ice with 30 s intervals for cooling). The obtained cellular extracts were centrifuged twice at 30,000g for 10 min to obtain clear supernatant. Ni-NTA resin (Qiagen, Venlo, The Netherlands) was washed 3 times and resuspended in 1 mL of extraction buffer before incubated with the centrifuged extracts at 4°C for 1 h with nutation. After removing the supernatant by centrifuging the slurry at 3,200g for 5 min, the resins were transferred to a gravity flow column (Bio-Rad Laboratories, Hercules, CA, USA) and washed with 3 column volumes of wash buffer (50 mM NaPO_4_, 300 mM NaCl, 40 mM imidazole, and 5 mM 2-mercaptoethanol, pH 8). Two milliliters of elution buffer (50 mM NaPO_4_, 300 mM NaCl, 3 M imidazole, and 5 mM 2-mercaptoethanol, pH 8) was added to the column and allowed to incubate with the resin for 1 min. The elutes were then diluted into 15 mL of storage buffer (extraction buffer without imidazole), concentrated using 10-kD centrifugal filter units (EMD Millipore, MA, USA) till 1,000-fold dilution. A final volume of 40% (v/v) glycerol-elution solution was prepared and stored at -20°C. Immunoblot with the anti-His-antibody conjugated to peroxidase (BMG-His-1 monoclonal antibody; Roche, Basel, Switzerland) was used to confirm the presence of enzymes.

### Transient expression and purification of SnASFF1 protein

ASFF1 protein expression and purification was carried out exactly as described before(*15*). SnASFF1 full-length open reading frame was amplified from *S. nigrum* young leaf and peduncle cDNA using c70979_Fw and c70979_Rv primer and cloned into the pEAQ-HT vector(*50*) using NruI-HF and SmaI restriction sites and 2× Gibson Assembly master mix (NEB, Ipswich, MA) according to the manufacturer’s instructions. The completed vector was subsequently transformed into *Agrobacterium tumefaciens* LBA4404 cells. Single colonies from LB agar plates with rifampicin (50 µg mL^-1^) and kanamycin (50 µg mL^-1^) were used to inoculate 50 mL of YEP medium with the same antibiotics in the same concentration. After overnight incubation at 28°C, 300 rpm, culture was harvested and washed in 50 ml of buffer A [10 mM 2-ethanesulfonic acid (MES; Sigma-Aldrich, St. Louis, MO, USA) at pH 5.6 and 10 mM MgCl_2_] before resuspended to a final OD600 = 1.0 with buffer A with 200 µM acetosyringone (Sigma-Aldrich, St. Louis, MO, USA). The suspension was incubated at room temperature with gentle rocking for 4 hours and then infiltrated into fully expanded leaves of 6-week-old *Nicotiana benthamiana* plants using a needleless 1-mL tuberculin syringe. Infiltrated leaves were harvested, deveined, and flash-frozen in liquid nitrogen after 7-8 days. Tissue was powdered under liquid nitrogen and added to 140 mL of ice-cold buffer B [25 mM 3-[4-(2-hydroxyethyl)piperazin-1-yl]propane-1-sulfonic acid (EPPS) at pH 8.0, 1.5 M NaCl, 1 mM EDTA with 2 mM dithiothreitol (DTT), 1 mM benzamidine, 0.1 mM phenylmethansesulfonyl fluoride, 10 µM trans-epoxysuccinyl-l-leucylamido(4-guanidino)butane (E-64), and 5% (w/v) polyvinylpolypyrrolidone (PVPP); all reagents were obtained from Sigma-Aldrich (St. Louis, MO, USA) except DTT, which was obtained from Roche Diagnostics (Risch-Rotkreuz, Switzerland)]. The mixture was stirred for 4 hours at 4°C before filtered through six layers of Miracloth and centrifuged at 27,000g at 4°C for 30 min. After passed through a 0.22-µm polyethersulfone filter (EMD Millipore, Billerica, MA), the supernatant was loaded onto a HisTrap HP 1-mL affinity column and eluted using a gradient of 10 to 500 mM imidazole in buffer B using an ÄKTA start FPLC module (GE Healthcare, Uppsala, Sweden). Fractions were analyzed by SDS–polyacrylamide gel electrophoresis, and the presence of SnASFF1-HT was confirmed by immunoblot using the BMG–His-1 monoclonal antibody (Roche, Basel, Switzerland). Purified SnASFF1 proteins were stored in 100 mM sodium acetate (pH 4.5) with 40% (v/v) glycerol at -20°C.

### Enzyme assays

Unless otherwise specified, all BAHD forward enzyme assays were performed by incubating purified recombinant proteins in 60 µL of 100 mM ammonium acetate (pH 6.0) buffer with 100 µM acyl-CoA and an acyl chain acceptor (unmodified sugar or purified or fractionated acylsugar acceptors in an ethanol:water mixture (1:1 v/v)) at 30°C for 30 min. After the incubation, 2 volumes of stop solution [acetonitrile:isopropanol (1:1) with 0.1% (v/v) formic acid and 1 µM telmisartan as internal standard] was added to the assays and mixed. Reactions were centrifuged at 17,000g for 5 min, and the supernatants were stored at -20°C after being transferred to LC-MS vials. SnASFF1 enzyme assays were carried out in a similar manner in the absence of acyl-CoAs. For BAHD reverse enzyme assays, 100 µM free Coenzyme A were used in place of acyl-CoAs. For negative controls, enzymes boiled at 95°C for 10 min were used in place of active enzymes. Methods used to determine the apparent *K*_m_ value for different substrates were performed as previously described(*11*). Tested sugars, free coenzyme A and nC10-, nC8-, iC5-, iC4-CoAs were obtained from Sigma-Aldrich, St. Louis, MO, USA, whereas iC10-CoA and aiC5-CoA were produced following method described in Kawaguchi et al., 1981(*51*).

### Purification of S1:10 and S2:14

Purifications were performed using a Waters 2795 Separations Module (Waters Corporation) and an Acclaim 120 C18 HPLC column (4.6 mm × 150 mm, 5 µm; Thermo Fisher Scientific, Waltham, MA, USA) with a column oven temperature of 30°C and flow rate of 1 mL/min. For S1:10 purification, the mobile phase consisted of water with 0.1% (solvent A) and acetonitrile (solvent B). For *in vivo* S2:14 purification, two purification methods where solvent B consist of acetonitrile and methanol, respectively, were used in a sequential manner to enhance purity of compounds. The HPLC methods used for separating S1:10 and S2:14 are described in Table S10. Fractions were collected using a 2211 Superrac fraction collector (LKB Bromma, Stockholm, Sweden).

### VIGS Analysis

VIGS target regions were selected to have a low chance of altering expression of trichome expressed non-target genes(*14*). These fragments were amplified from *S. nigrum* young leaf and peduncle cDNA using primers listed in Table S9 and were cloned into pTRV2-LIC as previously described (*16*). In short, the PstI-HF-linearized pTRV2-LIC vector and all PCR fragments were separately incubated in 5-µL T4 DNA polymerase reactions with 5 mM dATP or dTTP, respectively. The reactions were incubated at 22°C for 30 min, then 70°C for 20 min, and then stored on ice. Two microliters of desired PCR reaction were mixed with 1 µL vector reaction and incubated at 65°C for 2 min, followed by 22°C for 10 min. Sequenced constructs and pTRV1 were transformed into *A. tumefaciens* strain GV3101 using the protocol described previously(*15*).

The vacuum infiltration protocol was adapted from Hartl et al. (2008) (*52*). Twenty-five milliliters of LB medium with kanamycin (50 µg mL^-1^), rifampicin (50 µg mL^-1^), and gentamicin (10 µg mL^-1^) were inoculated with a single colony of the respective Agrobacterium strain from plate. After overnight incubation at 28°C and 225 rpm, cells were harvested by centrifugation at 3,200g for 10 min and resuspended to a final OD600 = 1 with the induction media (10 mM MES, pH 5.6, and 10 mM MgCl_2_). An equal volume of pTRV1 suspension was mixed with different pTRV2-LIC constructs suspension. 200 µM acetosyringone was added to each mixture before the suspension was incubated in the dark at room temperature with gentle rocking for 3 hours. Seven to ten days-old *S. nigrum* seedlings were carefully transferred from the petri dishes to the bacteria solution. Vacuum was applied in a desiccator for 2 min, followed by slow release of the vacuum. Infiltrated seedlings were kept for 3 d under indirect light, then planted in Jiffy-7® peat pellets and grown in a climate chamber as described above. Metabolite and tissue samples were harvested approximately 3 weeks post infiltration and phytoene desaturase silencing efficiency was monitored. The QuanLynx function in MassLynx v4.1 (Waters Corporation, MA, USA) was used to integrate extracted ion chromatograms from untargeted LC-MS data as described previously (*29*). In short, all quantifications were performed using extracted ion chromatograms of the *m/z* value for the relevant [M+NH_4_]^+^ or [M+HCOO]^-^ adduct ions using a mass window of *m/z* 0.05. Isomeric forms of acylsugars (including anomers) were quantified in corresponding groups as shown in Table S1. The retention time window was adjusted for each compound to include all isomers. Peak area of telmisartan was quantified and used as an internal reference. Then, acylsugar quantities (per mg) were calculated by normalizing peak areas to the internal standard peak area and dry leaf weight, whereas acylsugar ratios were calculated by comparing peak areas over internal standard peak area of acylsugars of interest.

### qPCR Analysis

RNA was extracted with the RNeasy Plant Mini Kit including on-column DNase digestion (Qiagen, Venlo, The Netherlands), according to the manufacturer’s instructions. RNA was quantified with a Nanodrop 2000c instrument (Thermo Fisher Scientific, Waltham, MA, USA). cDNA was synthesized using 1mg of the isolated RNA and SuperScript II Reverse Transcriptase (Invitrogen, Carlsbad, CA, USA). The cDNA samples were diluted 40-fold (10-fold initial dilution and 4-fold dilution into qPCRs). qPCRs (10µL) were created with SYBR Green PCR Master Mix (Thermo Fisher Scientific, Waltham, MA, USA), and primers were used at a final concentration of 200 nM. RT_SnASAT1_F/R, RT_SnASAT2_F/R, RT_SnASFF1_F/R, RT_SnAGAT1_F/R, RT_Actin_1_F/R, and RT_Actin_3_F/R primers were used to detect *SnASAT1*, *SnASAT2*, *SnASFF1*, *SnAGAT1*, *ACTIN1*, and *ACTIN3* transcripts, respectively (Table S9). Reactions were carried out with a QuantStudio 7 Flex Real-Time PCR System (Applied Bio-systems) by the Michigan State University RTSF Genomics Core. The following temperature cycling conditions were applied: 50°C for 2 min, 95°C for 10 min, and 40 cycles of 95°C for 15 s and 60°C for 1 min. Relative expression of *SnASAT1*, *SnASAT2*, *SnASFF1*, and *SnAGAT1* was calculated with the ΔΔCt method (*53*) and normalized to the geometric mean of *ACTIN1* and *ACTIN3* transcript levels. The mean expression values of the transcripts in the control plants were used for normalization. Three to four technical replicates were used for all the qPCRs.

### Statistical Analysis

All statistical analyses were performed using the “stats” R package (R Core Team, 2017). Welch two-sample *t* tests were executed on metabolites and transcript abundance data using the “t.test” command. The power of these analyses was determined using the “power.t.test” function.

## Supporting information

All Supplementary Tables

## Acknowledgments

We acknowledge the MSU RTSF Mass Spectrometry and Metabolomics Core Facilities for their support with LC-MS analysis. We thank Dr. D. Holms and L. Xie at the Michigan State University Max T. Rogers NMR Facility for technical assistance. We thank Dr. Dan Lybrand for helpful advice and generously providing the pET28b-SnASAT1 construct. We also acknowledge the valuable feedback received from members of the Last lab.

## Author contributions

Design: YRL, TMA, ADJ, RLL

NMR experiments and data analysis: TMA, PDF, ADJ

Biochemical and biological experiments and data analysis: YRL, PDF, REA and EMC Interpretation and preparation of figures and table: YRL, TMA, PDF

Writing – original draft: YRL, RLL

Writing – review and editing: YRL, TMA, PDF, REA, EMC, ADJ and RLL

## Data and materials availability

All data needed to evaluate the conclusions in the paper are present in the paper and/or the Supplementary Materials. Additional data related to this paper may be requested from the authors. The constructs pET28b-SnASAT1, pET28b- SnASAT2, pET28b-SnAGAT1, and pEAQ-HT-SnASFF1 can be provided by Y.-R.L pending a completed material transfer agreement. Requests for biological materials or data should be submitted to R.L.L. at.

## Supplementary Text

### Acylsugar annotation

Acylsugar structures were inferred from positive and negative mode MS collision-induced dissociation (CID) as previously described (*15, 29, 32, 34*). In short, we employed two discrete negative mode CIDs, where the lower energy (15V) removes acyl chains from acylsugars in a stepwise manner and the higher energy (30V) creates fatty acid fragment ions from acylsugar acyl chains. For acylsucroses, the broken glycosidic linkage by positive mode CID (15 or 30V) provided additional information about acyl chain attachment: a neutral loss of the furanose ring resulting in [M+NH_4_-FRUC]^+^ accumulation would indicate that all acyl chains are attached to the glucopyranose ring. Ring acylation information is indicated in Table S11. Examples of typical fragmentation of *S. nigrum* di- and triacylglucoses and triacylinositols are depicted in Fig. S2, whereas examples of mass spectra of the mono-and di-acylsucrose intermediates on acylglucose biosynthetic pathways are depicted in Fig. S7C and S9.

Below is the stepwise protocol with a list of criteria used in this manuscript to annotate and report acylsugars with different confidence levels. Putative acylhexoses that meet all the following criteria A-F were annotated with highest confidence in this manuscript. Those not meeting criteria E and/or F were reported with medium confidence, whereas potential acylhexoses that only meet criteria A were reported with low confidence. Confidence levels for each reported acylhexose are specified in Table S11.

(A) **Exact mass.** Experimental exact masses of potential acylsugars (*m/z* listed in Table S11) were compared with theoretical exact masses of possible acylsugars. A mass difference within 10 ppm met this criterion. Then, we generated extracted ion chromatograms (EICs) of each potential acylsugar to compare with the EICs generated in B and C.
(B) **Presence of co-eluting sugar core fragments.** For acylglucoses, EICs of characteristic glucose core structure (*m/z* 127.04 in 15 or 30V positive mode CID and *m/z* 143.03 in 0V negative mode CID) were generated. For acylsucroses, EICs for the sugar core structure on acylsucroses (*m/z* 341.10 and 323.09 in negative mode) were generated. As acylinositols fragment less well in negative mode, the characteristic *myo*-inositol core fragment ion (*m/z* 127.04) was used to generate EICs in positive mode CID (30V).
(C) **Presence of co-eluting fatty acid carboxylate fragments in high CID potential (30V).** EICs of the carboxylate fragments of C4 (*m/z* 87.04), C5 (*m/z* 101.06), unsaturated C5 (*m/z* 99.05), C6 (*m/z* 115.07), C7 (*m/z* 129.09), C8 (*m/z* 143.10), C9 (*m/z* 157.12), C10 (*m/z* 171.14), C11 (*m/z* 185.15) and C12 (*m/z* 199.17) were generated in highest CID potential (30V) in negative mode.
(D) **Presence of co-eluting fragments that correspond to stepwise loss of acyl chains**. The EICs generated in (A), (B) and (C) were closely examined for co-eluting peaks that indicate potential acylsugars. For each potential acylsugar, spectra from the top of the peak were then carefully examined for stepwise losses of acyl chains in negative mode CID at 15V. Acylinositol fragmentation for this criterion was observed in positive mode due to poor negative mode fragmentation^5,6^.
(E) **Confirmed by MS/MS.** The above annotation were then confirmed with data-dependent (DDA) MS/MS when applicable.
(F) **Consistency across samples.** Acylsugar presence was monitored across multiple metabolite extracts. Potential acylsugars met this criterion when they were detected across all samples analyzed.

As an example, the three peaks with *m/z* 491.25 with retention time 52.66, 53.22 and 54.02 min on the 110-min share identical (δ_m_ = 0.1 ppm) with the [M+formate]^-^ of H3:16 (Fig. S3A). To confidently annotate these peaks, co-elution of [M+formate]^-^ (*m/z* 491.25) and glucose fragment ion (*m/z* 143.03) was verified on EICs generated in 0V negative mode. Co-eluting carboxylate anion were identified for C10 carboxylate anion (*m/z* 171.14) and C4 carboxylate anion (*m/z* 87.04) in 30V negative mode. Presence of C4 and C10 acyl chains was further confirmed by ion abundance of [M-C4-H_2_O]^-^ (*m/z* 357.19) and [M-C4-C10-2H_2_0]^-^ (*m/z* 185.04) on CID at 15V negative mode. As these fragmentation were confirmed on MS/MS fragmentation of [M+formate]^-^ (*m/z* 491.25), we report the three peaks between 52-55 min with *m/z* 491.25 as triacylhexoses H3:16(2,4,10) with high confidence level based on LC-MS data.

**Fig. S1.**
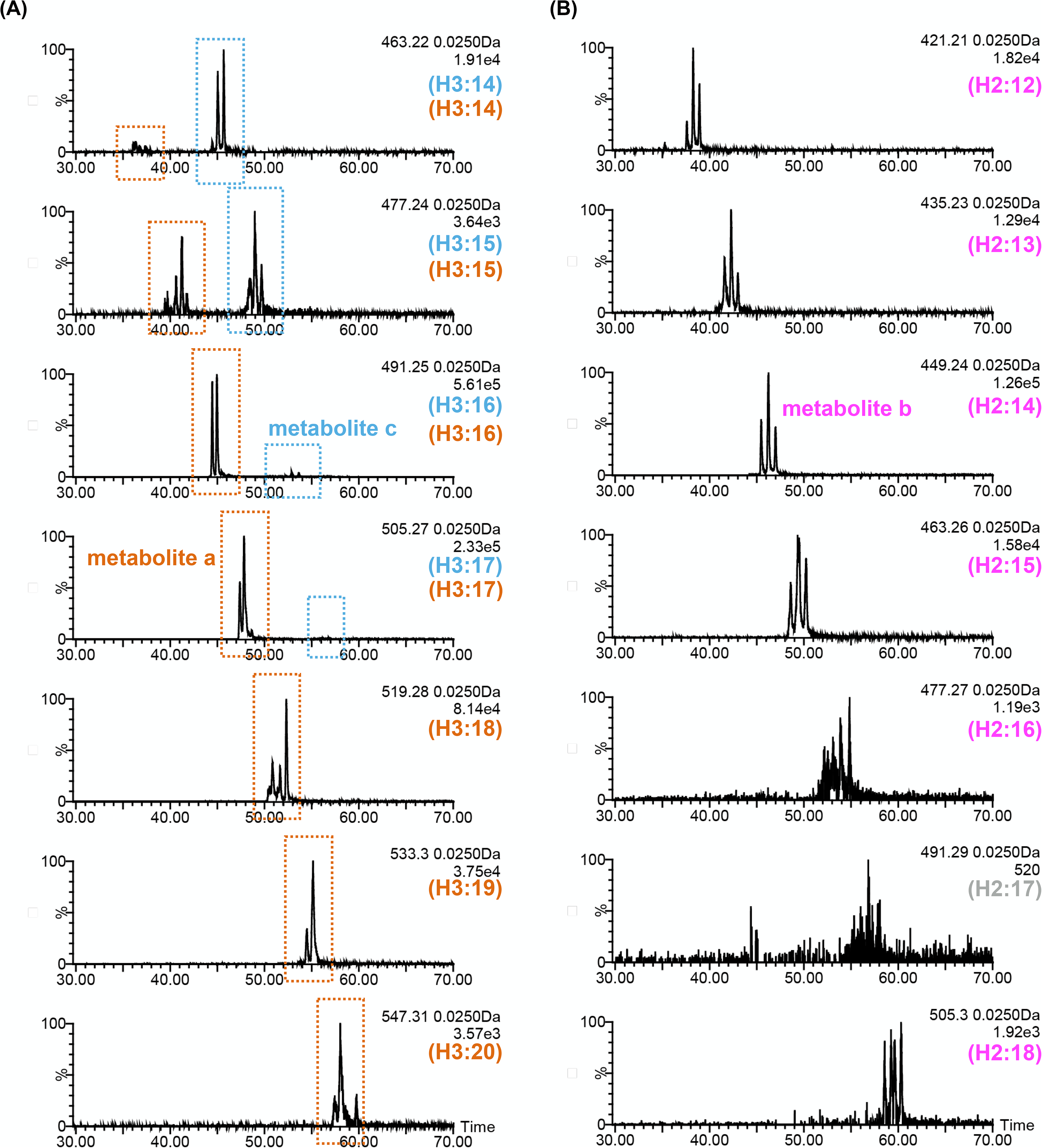
Chromatographs of abundant acylhexoses in *S. nigrum*. **(A)** Extracted ion chromatograms (EICs) from negative ion mode LC-MS analyses showing selected *m/z* values correspond to [M+formate]^-^ of hexose triesters. Observed peaks in these chromatograms separate into two distinct retention time groups: peaks corresponding to triacylglucoses, such as metabolite c, are highlighted with cyan dash boxes, and those corresponding to triacylinositols, such as metabolite a, are highlighted in orange dash boxes. **(B)** Extracted ion chromatograms from negative ion mode LC-MS analyses showing selected *m/z* values correspond to [M+formate]^-^ of hexose diesters. Numerous instances suggested common chromatographic overlap of isomeric acylhexoses was common. The corresponding *m/z* of these compounds are listed in Table S1 and Supplementary Text.

**Fig. S2.**
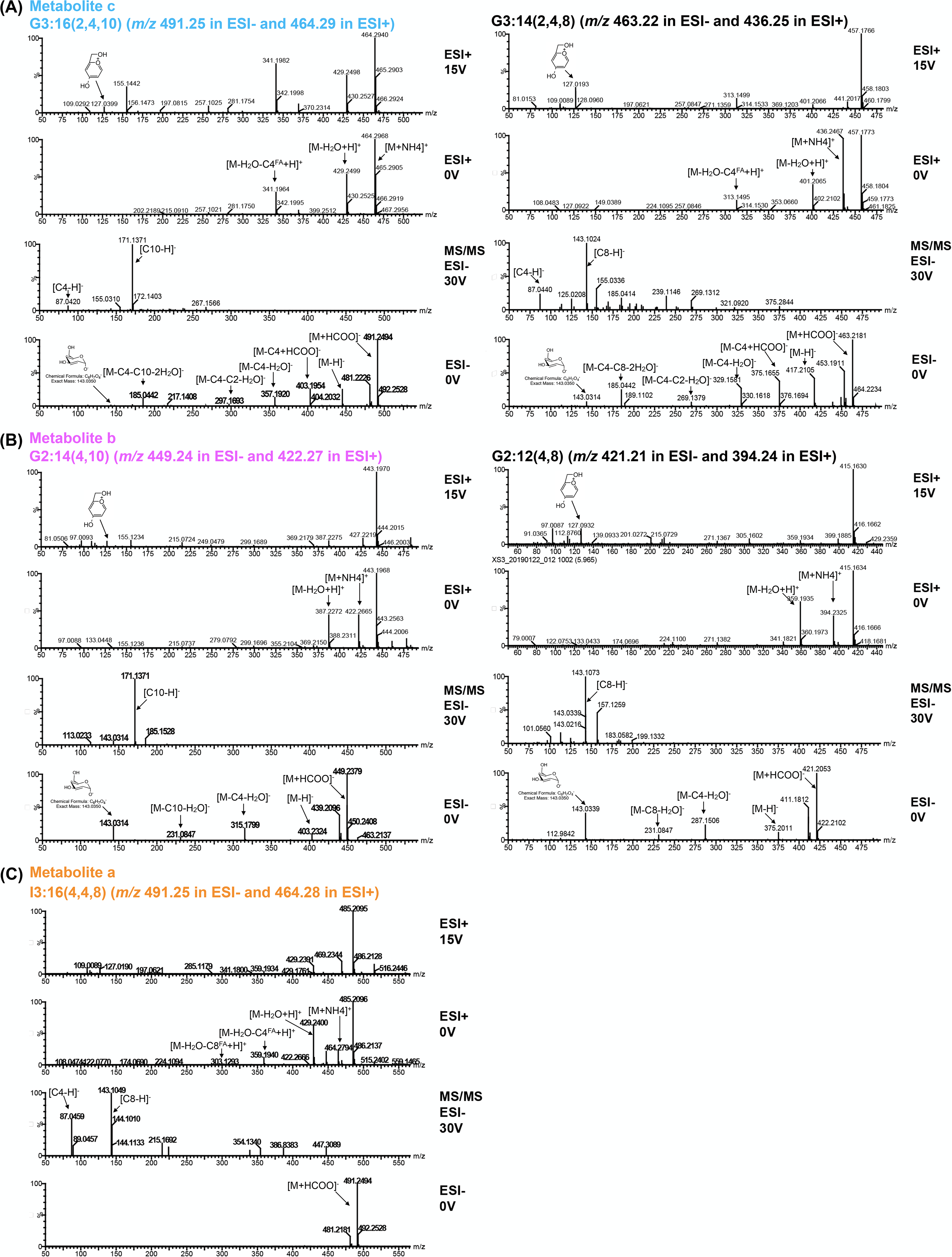
Mass spectra of abundant acylhexoses in *S. nigrum*. *S. nigrum* di- and triacylglucoses and triacylinositols fragment in either negative or positive ion mode by losing the short and medium-length acyl chains as neutral fatty acids, which can also be seen in negative ion mode. These observations allow for determination of the number and length of acyl chains attached to each acylglucose. For more detail, see Supplementary Text. **(A)** Fragmentation of *S. nigrum* triacylglucoses G3:16 and G3:14. Fragmentation of G3:16 in 0V ESI- mode is characterized by the loss of C4, C10 and C2 ketenes. ESI-MS/MS spectrum (30V) generated from [M+formate]^-^ of G3:16 reveals the presence of C4 (*m/z* 87.04) and C10 (*m/z* 171.14) fatty acids. Similarly, fragmentation of G3:14 in ESI- mode is characterized by the loss of C4, C8 and C2 ketenes under no collision energy (0V), while 30V ESI-MS/MS of product ions from [M+formate]^-^ of G3:14 reveals the presence of C4 (*m/z* 87.04) and C8 (*m/z* 143.10) fatty acids. Fragmentation of both triacylglucoses in ESI+ mode (0 and 15V) results in the loss of a C4 fatty acid, followed by loss of a C10 or C8 fatty acid, respectively. **(B)** Fragmentation of *S. nigrum* diacylglucoses G2:14 and G2:12. Fragmentation of G2:14 in ESI- mode is characterized by the loss of C4 and C10 ketenes under no collision energy (0V). ESI- MS/MS spectra (30V) of product ions generated from [M+formate]^-^ of G2:14 reveals the presence of C4 (*m/z* 87.04) and C10 (*m/z* 171.14). Similarly, fragmentation of G2:14 in ESI- mode is characterized by the loss of C4 and C8 ketenes under no collision energy (0V), while 30V ESI- MS/MS of product ions from [M+formate]^-^ of G2:12 reveals the presence of C4 (*m/z* 87.04) and C8 (*m/z* 143.10). ESI+ mode (15V) of both diacylglucoses reveal the loss of C4 and a C10 or C8, respectively, while 0V ESI+ results in little fragmentation. **(C)** Fragmentation of I3:16 from *S. nigrum*. Fragmentation of I3:16 in ESI+ mode is characterized by the loss of C4 and C10 fatty acids under no collision energy (0V). ESI- MS/MS spectra (30V) of product ions generated from [M+formate]^-^ of I3:16 reveals the presence of C4 (*m/z* 87.04) and C8 (*m/z* 143.10). In contrast to acylglucoses, 0V ESI- result in little fragmentation for acylinositols.

**Fig. S3.**
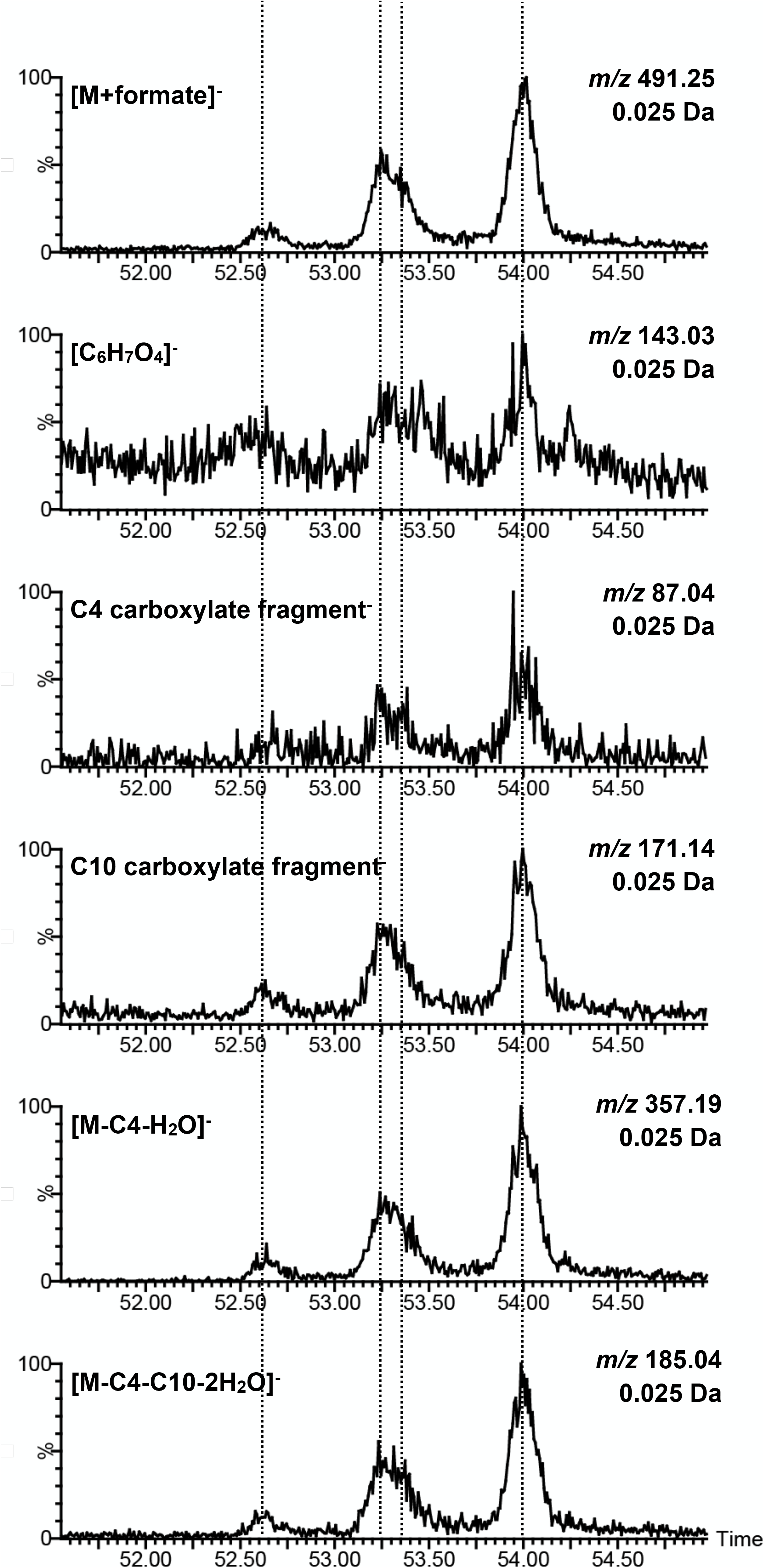
Co-chromatography of metabolite c and characteristic fragment ions under different collision-induced dissociations (CID). Extracted ion chromatograms (EICs) of (from top to bottom) [M+formate]^-^ of metabolite c (*m/z* 491.25) in 0V ESI-, glucose core structure [C_6_H_7_O_4_]^-^ (*m/z* 143.03) in 0V ESI-, C4 (*m/z* 87.04) and C10 (*m/z* 171.14) carboxylate fragment under CID potential 30V, and [M-C4-H_2_O]^-^ (*m/z* 357.19) and [M-C4-C10-H_2_O]^-^ (*m/z* 185.04) fragment ions under CID potential 15V. For more details, see Supplementary Text.

**Fig. S4.**
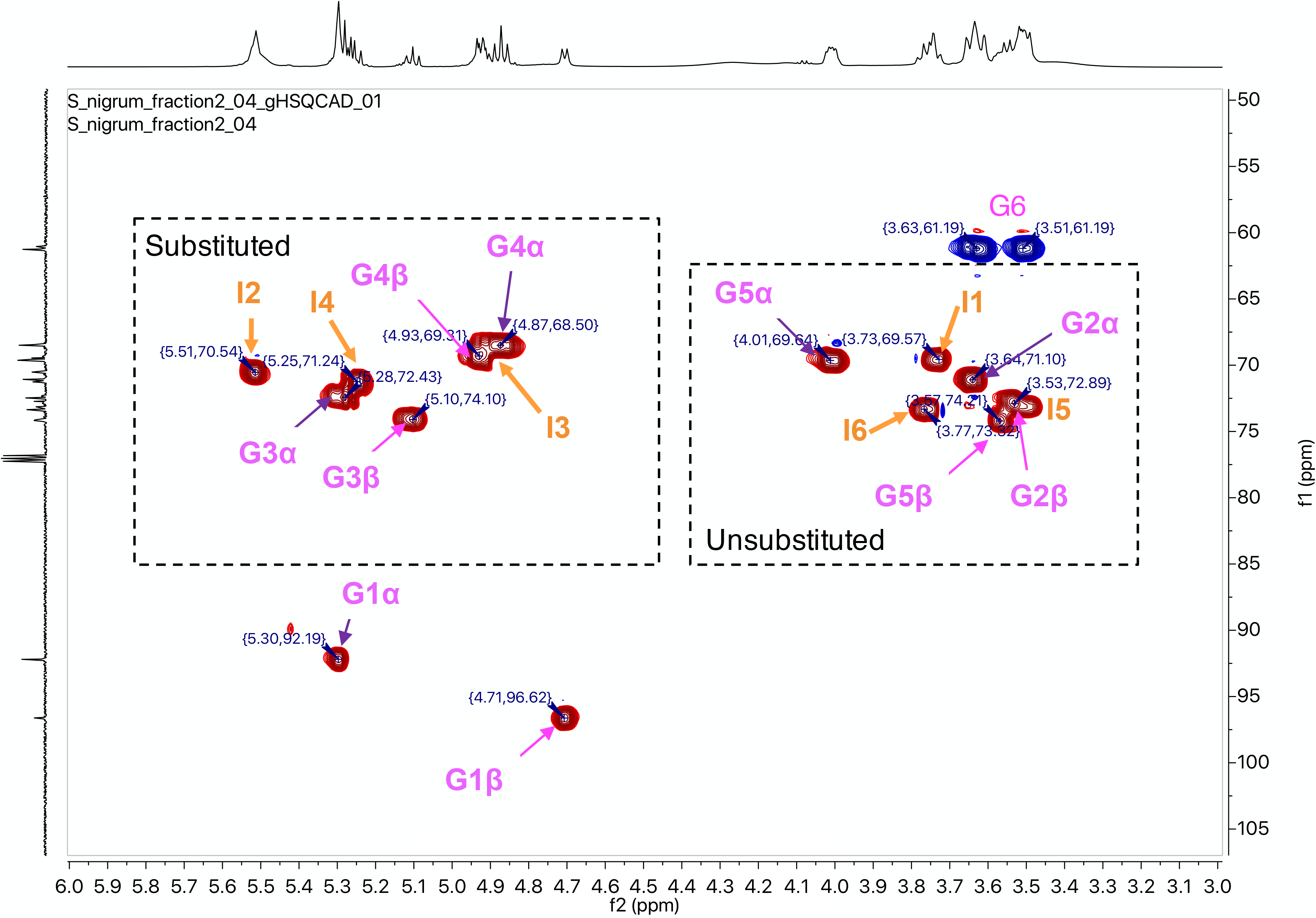
HSQC spectrum reveals two type of acylsugars in the more-polar fraction of *S. nigrum* leaf surface extract. HSQC spectrum of fraction 2 shows signals from diacylglucoses (magenta) and triacylinositols (orange) that were confirmed after HSQC-TOCSY separating correlation signals.

**Fig. S5.**
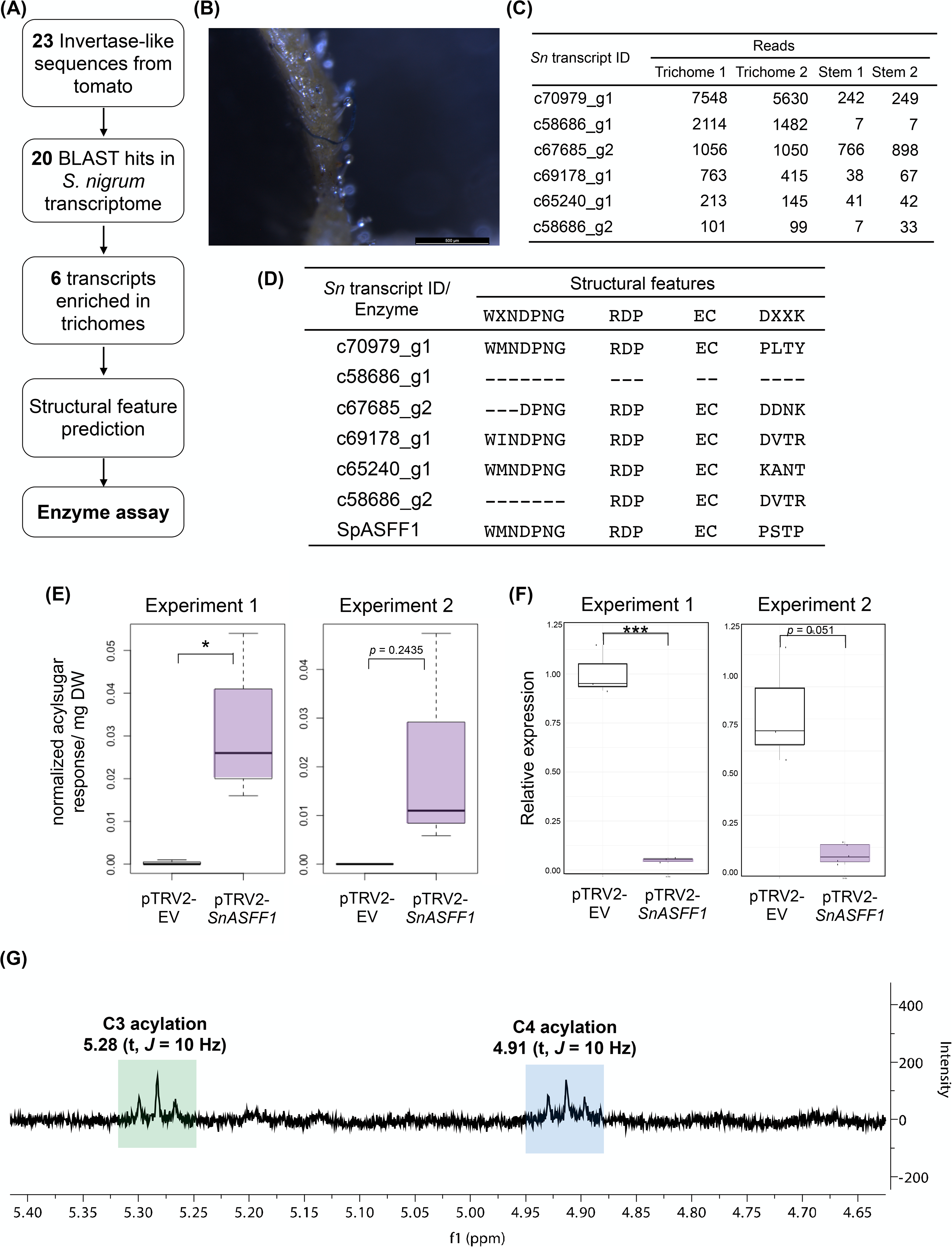
Identification of a trichome-expressed ASFF candidate involved in *S. nigrum* acylglucose biosynthesis. **(A)** Flow chart of identifying ASFF candidate transcripts in *S. nigrum* transcriptome. **(B)** Close-up photo of a *S. nigrum* peduncle with acylsugar droplets accumulating on trichome tips. Excessive acylsugars produced on the peduncle appear to drip down and coat the trichome-free fruit. **(C)** Six invertase-like candidate transcripts are enriched in *S. nigrum* trichome RNA-Seq data from Moghe et al. (2017) **(D)** Structural features identified a transcript, *c70979_g1 (SnASFF1)*, with conserved catalytic sites and a non-canonical substrate binding site, as seen on SpASFF1. **(E)** Comparison of S2:14(4,10) accumulation in *SnASFF1*-targeted and empty vector VIGS plants from two independent experiments. Acylsugars were analyzed using LC-MS in ESI- mode. Peak areas of S2:14(4,10) (*m/z* 611.29) were integrated under negative mode and normalized to the internal standard telmisartan and dry leaf weights. See Table S5 for details. **(F)** qPCR analysis of *SnASFF1* transcript abundance in VIGS plants from two independent experiments. Relative transcript abundance was generated using the ΔΔCt method (*53*) and normalized to the geometric mean of two actin gene transcript levels. Significant levels are shown (*, *p* < 0.05; **, *p* < 0.01; ***, *p* < 0.001; Welch’s two sample t-test). **(G)** ^1^H NMR of S2:14(4,10) purified from *SnASFF1*-targeted VIGS plants. Two triplet peaks at 5.28 (green box) and 4.91 ppm (blue box) indicate the 3- and 4-positions on sucrose, respectively, are acylated.

**Fig. S6.**
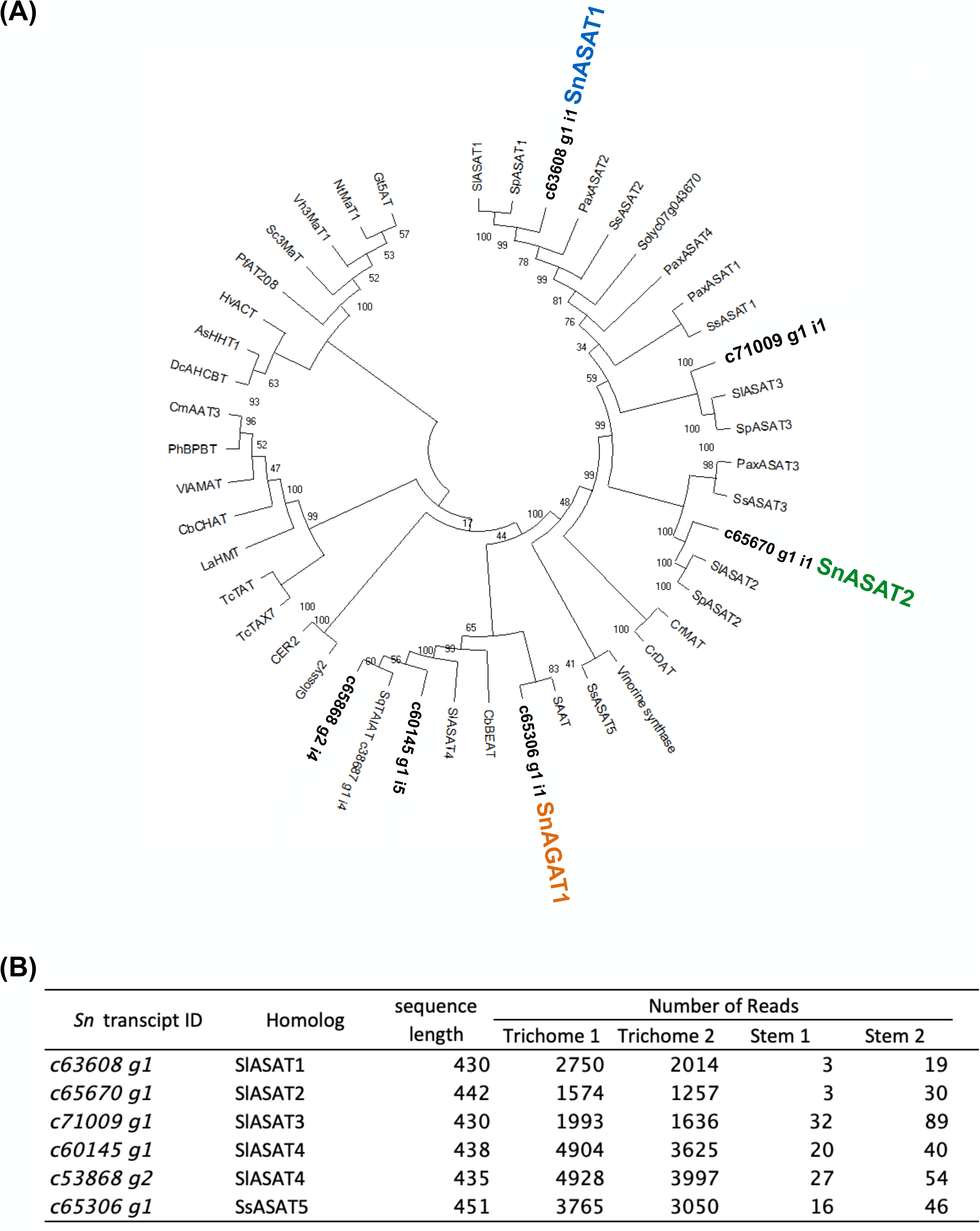
Six ASAT homologs are preferentially expressed in *S. nigrum* trichomes. **(A)** Phylogenetic analysis of ASAT candidates and previously characterized BAHDs. The phylogenetic tree shows several previously characterized ASATs and other BAHD acyltransferases. BAHD candidates in S. nigrum are marked in bold, whereas characterized SnASATs are marked with respective colors (SnASAT1 in blue, SnASAT2 in green and SnAGAT1 in orange). **(B)** Sequence length and RNA-Seq data for homologs of characterized ASATs in *S. nigrum*. RNA-Seq data are derived from Moghe et al. (2017).

**Fig. S7.**
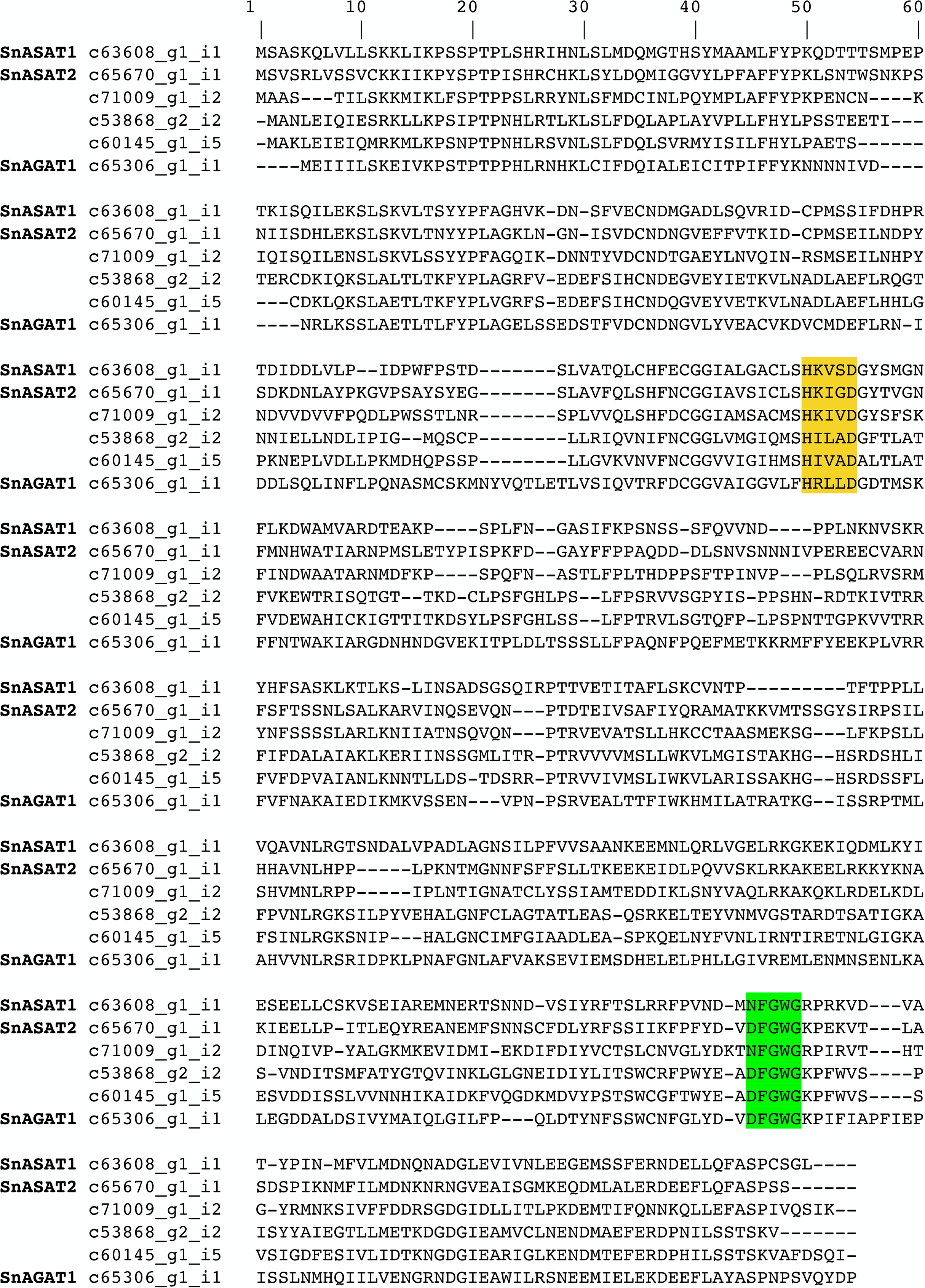
Multiple sequence alignment of candidate sequences. BAHD characteristic HXXXD motifs (yellow) and DFGWG-like motif (green) are highlighted. Hyphens indicate gaps in protein sequence.

**Fig. S8.**
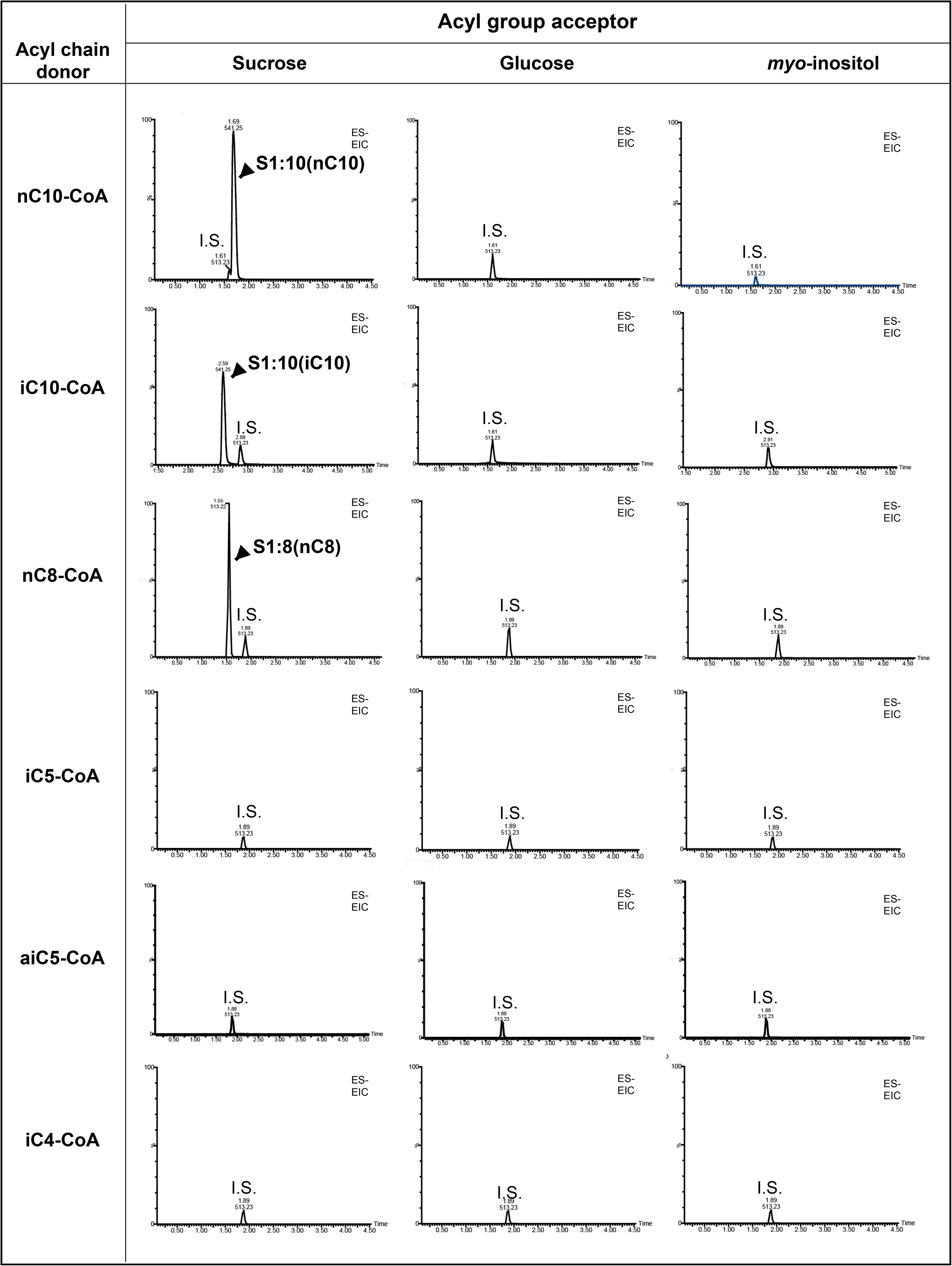
SnASAT1 activities with different acyl chain donors and acyl group acceptors. Acylsugars were analyzed using LC-MS in ESI- mode. Combined extracted ion chromatogram showing internal standard telmisartan (*m/z* 513.23) and expected products as formate adducts: top two rows from left to right, S1:10 (*m/z* 541.25), G1:10 (*m/z* 379.21) and I1:10 (*m/z* 379.21); third row from left to right, S1:8 (*m/z* 513.22), G1:8 (*m/z* 351.17) and I1:8 (*m/z* 351.17); fourth and fifth rows from left to right, S1:5 (*m/z* 471.18), G1:5 (*m/z* 309.12) and I1:5 (*m/z* 309.12); bottom from left to right, S1:4 (*m/z* 457.16), G1:4 (*m/z* 295.11) and I1:4 (*m/z* 295.11).

**Fig. S9.**
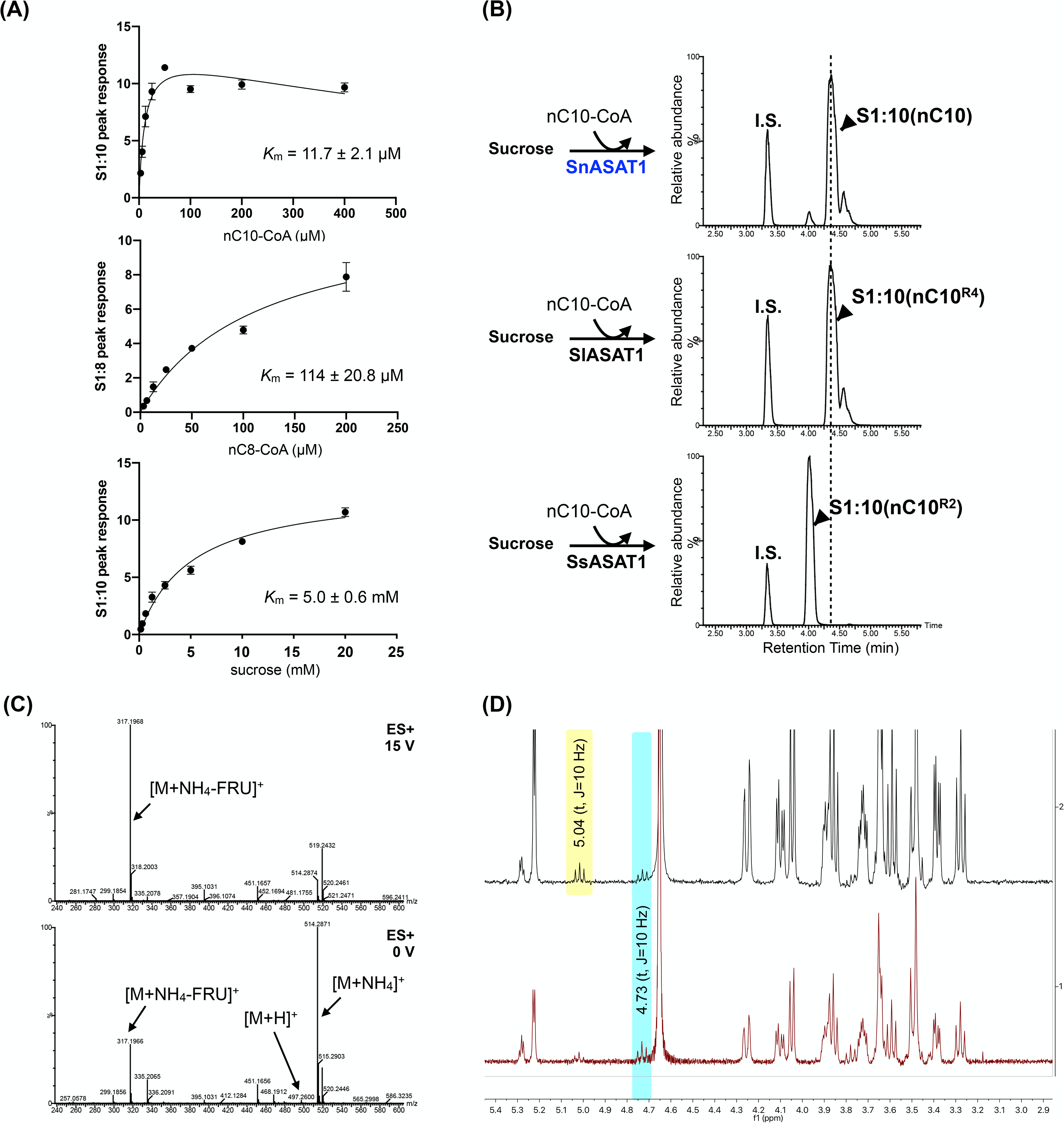
SnASAT1 generates monoacylsucroses with a medium-length acyl chain on the R_4_ position. **(A)** Kinetics analysis of different substrates for SnASAT1. Apparent *K*_m_ were calculated using the non-regression model in the GraphPad Prism 5 software. The enzyme assay products’ peak area was normalized to the internal standard telmisartan peak area and plotted for each concentration of the varying substrate. Error bars indicate standard error with n = 3. **(B)** Co-chromatography of S1:10(10) produced by SnASAT1 (top), SlASAT1 (middle) and SsASAT1 (bottom). Acylation positions of SlASAT1- and SsASAT1-produced S1:10(10) were verified by NMR^1,2^. **(C)** ESI+ spectra under CID potential 15V of product ions generated from [M+NH_4_]^+^ of SnASAT1-produced S1:10 results in neutral loss of unacylated fructose moiety. **(D)** Proton NMR determined the major S1:10(10) isomer is acylated at the R_4_ position [4(CH) = 4.73 (**t**, J=10Hz); top and bottom spectrum]. A second, chromatographically separable S1:10 minor isomer with acylation at R_6_ position [6(CH) = 5.04 (**t**, J=10Hz); top spectrum] accumulates after extended enzyme assay incubation time.

**Fig. S10.**
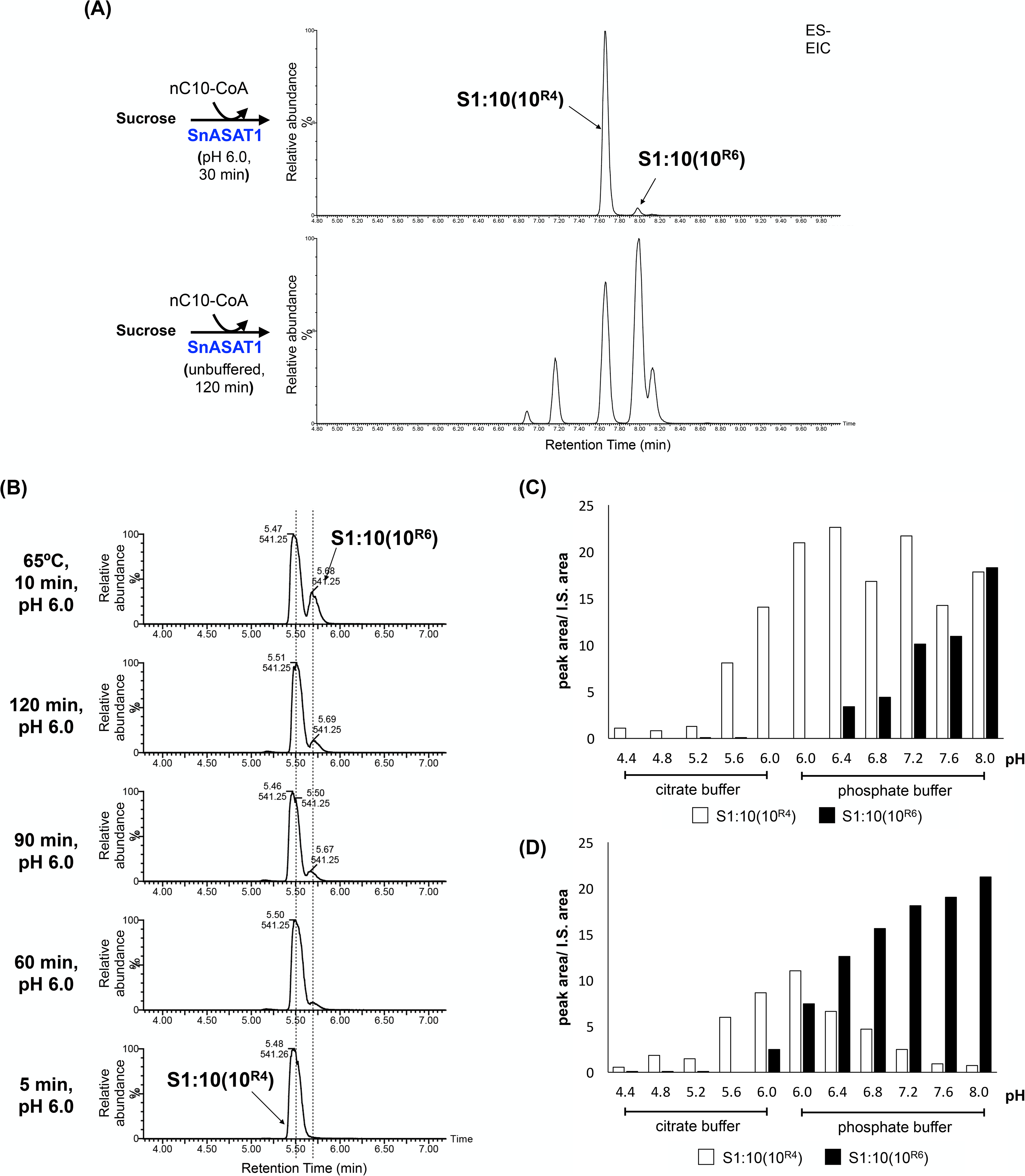
R_6_-acylated S1:10 isomer accumulates in SnASAT1 enzyme assays. **(A)** The co-chromatography of S1:10(10) isomers produced by SnASAT1 under pH 6.0 for 30 min (top) and unbuffered aqueous solution for 120 min (bottom). **(B)** LC-MS profile of enzyme assays carried out with different incubation conditions R_6_-acylated S1:10 isomer accumulation increases after extended enzyme assay incubation time (bottom four) and brief exposure to elevated temperature (65°C) (top). **(C)** The concentration of the R_6_-acylated isomer increases when enzyme assays were carried out in neutral-to-alkaline pH conditions, or **(D)** subjected to brief exposure to elevated temperature (65°C). Acylsugars were analyzed using LC-MS in ESI- mode. Extracted ion chromatograms are showing formate adducts of S1:10(10) (*m/z* 541.25). Peak areas of S1:10(10) were integrated under negative mode and normalized to the internal standard telmisartan.

**Fig. S11.**
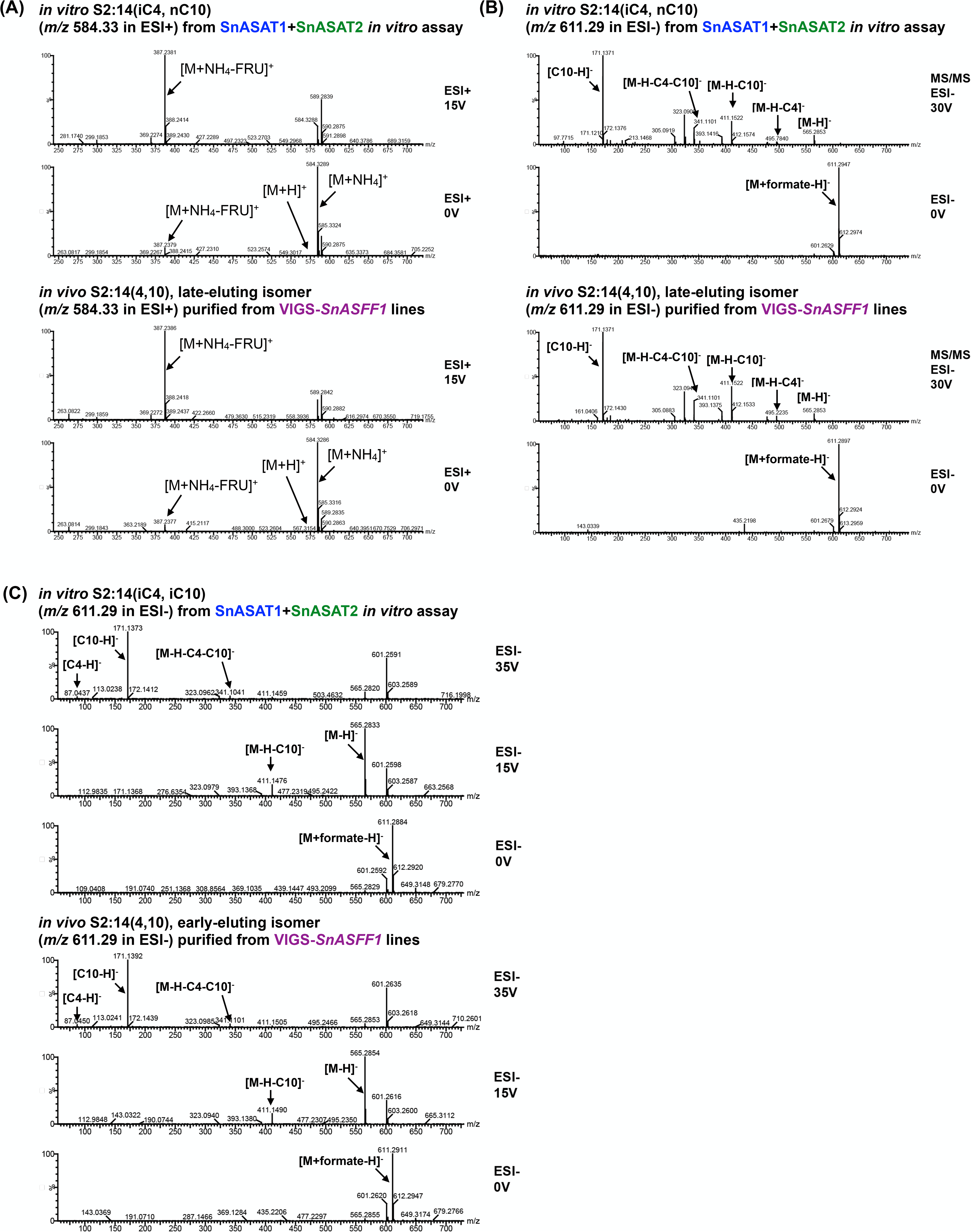
Mass spectra of the co-eluting *in vivo* and *in vitro* S2:14 isomers. **(A)** ESI+ mode fragmentation of the co-eluting *in vitro* S2:14(iC4, nC10) and *in vivo* late-eluting S2:14(4,10) results in the cleavage of the glycosidic linkage. Fragment analysis of the acylsucroses using collision-induced dissociation reveals a fragment ion (*m/z* 387.24) consistent with the furanose ring of sucrose conjugated to a C10 acyl chain, indicating that both acylsugars possess a C10 acyl chain on the furanose ring. **(B)** ESI- MS/MS spectra (30V) of product ions generated from [M+formate]^-^ of the co-eluting *in vitro* S2:14(iC4, nC10) and *in vivo* late-eluting S2:14(4,10) isomer are both characterized by the loss of C4 and C10 ketenes and the presence of C10 fatty acid anions (*m/z* 171.14). **(C)** Comparable fragmentation of the co-eluting *in vitro* S2:14(iC4, iC10) and *in vivo* early-eluting S2:14(4,10) isomer in ESI- mode reveals the loss of C4 and C10 ketenes under high collision energy (35V). The acyl chain combination is validated by the presence of C4 (*m/z* 87.04) and C10 fatty acid anions (*m/z* 171.14).

**Fig. S12.**
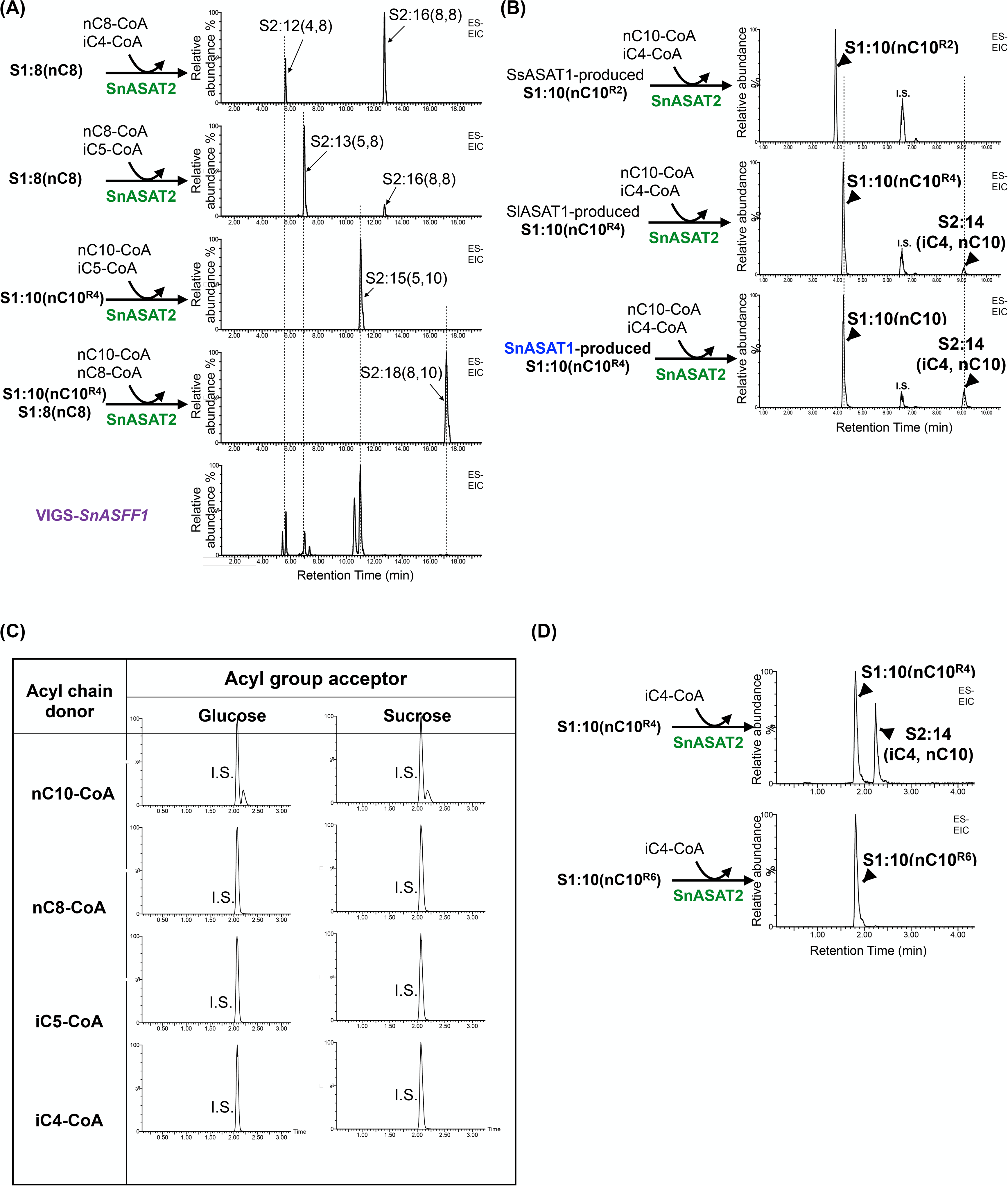
SnASAT2 activity with different acyl chain donors and acyl group acceptors. **(A)** LC-MS analysis of *in vitro* enzyme assay products indicates SnASAT2 takes SnASAT1-produced monoacylsucroses and iC4-, iC5 and nC8-CoAs as substrates to produce *in vitro* diacylsucroses (top four) that co-elute with *in vivo* diacylsucroses accumulating in VIGS-*SnASFF1* plants (bottom). All acylsugars were analyzed using LC-MS in ESI- mode. Combined extracted ion chromatogram showing enzyme assay products as formate adducts: S2:12 (*m/z* 583.26), S2:13 (*m/z* 597.28), S2:14 (*m/z* 611.29), S2:15 (*m/z* 625.31), S2:16 (*m/z* 639.33), S2:18 (*m/z* 667.36). **(B, C, D)** LC-MS analysis of *in vitro* enzyme assay products indicates SnASAT2 takes R_4_-acylated S1:10(10) as substrates, but not R_2_- or R_6_-acylated S1:10(10), unmodified glucose or unmodified sucrose. The S1:10 isomers used in panel D were purified from SnASAT1 enzyme assays and verified by NMR (Fig. S7D). All acylsugars were analyzed using LC-MS in ESI- mode. Combined extracted ion chromatogram showing internal standard telmisartan (*m/z* 513.23) and formate adducts of S1:10 (*m/z* 541.25) and S2:14 (*m/z* 611.29) in panel B and D, and S1:10 (*m/z* 541.25), G1:10 (*m/z* 379.21), S1:8 (*m/z* 513.22), G1:8 (*m/z* 351.17), S1:5 (*m/z* 471.18), G1:5 (*m/z* 309.12), S1:4 (*m/z* 457.16) and G1:4 (*m/z* 295.11) in panel C.

**Fig. S13.**
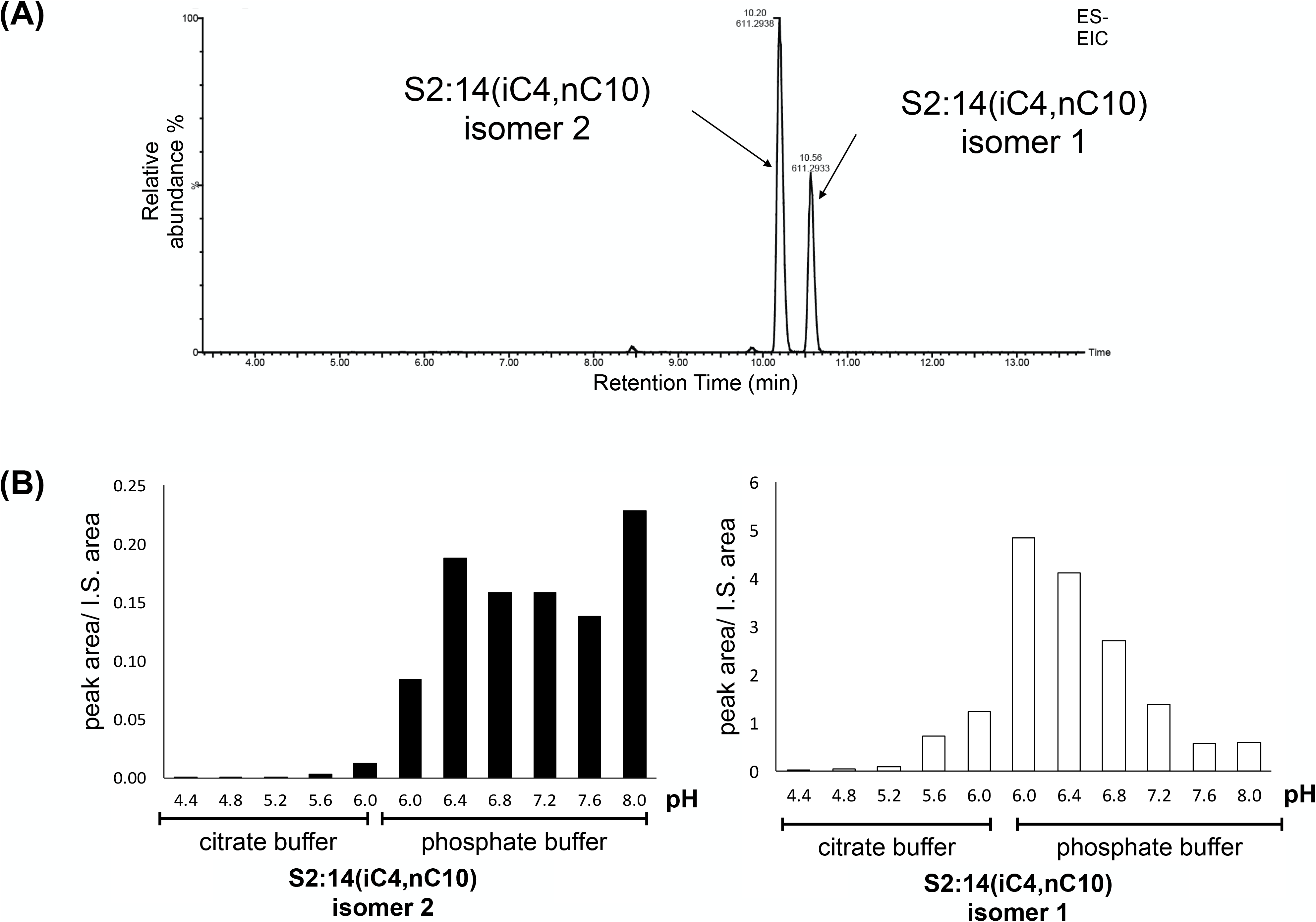
A second S2:14 isomer accumulates in SnASAT2 enzyme assays. **(A)** LC-MS analysis of SnASAT2 *in vitro* assay products showed a second isomer (isomer 2) accumulating in unbuffered water based solution for 120 min. **(B)** The concentration of the second isomer increases when enzyme assays were carried out in neutral-to-alkaline pH conditions. Acylsugars were analyzed using LC-MS in ESI- mode. Extracted ion chromatogram showing formate adducts of S2:14 (*m/z* 611.29). Peak areas of S2:14 were integrated under negative mode and normalized to the internal standard telmisartan.

**Fig. S14.**
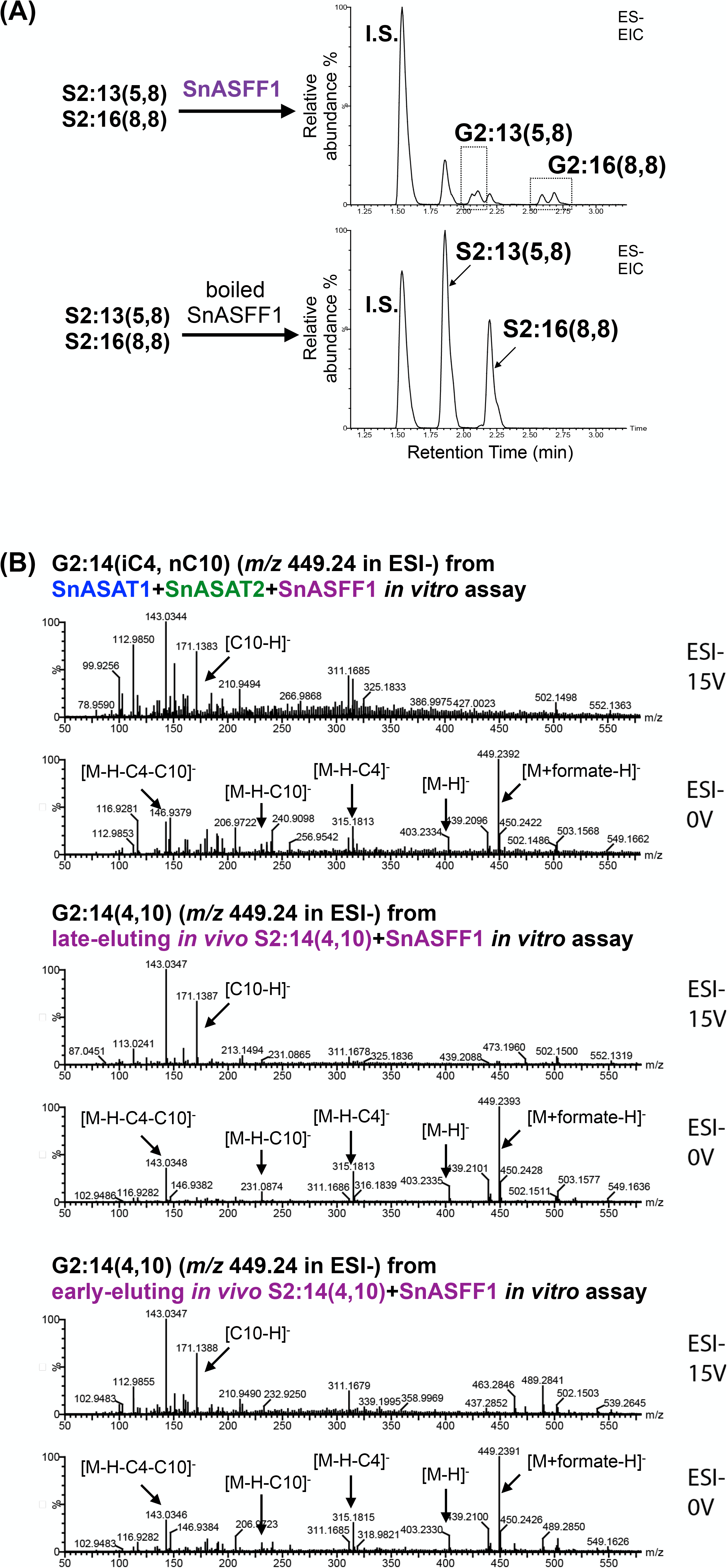
*In vitro* validation of involvement of SnASFF1 in *S. nigrum* acylglucose biosynthesis. **(A)** ESI- mode LC-MS analysis of *in vitro* enzyme assay products indicates SnASFF1 hydrolyzes S2:13(5,8) and S2:16(8,8) to produce G2:13(5,8) and S2:16(8,8). **(B)** The G2:14 isomers produced from *in vitro* (top) and *in vivo* (bottom two) S2:14 isomers generate comparable mass spectra that reveals the loss of C4 and C10 ketenes under no collision energy (0V) and the presence of C10 fatty acid anions (*m/z* 171.14) at elevated collision energy (15V).

**Fig. S15.**
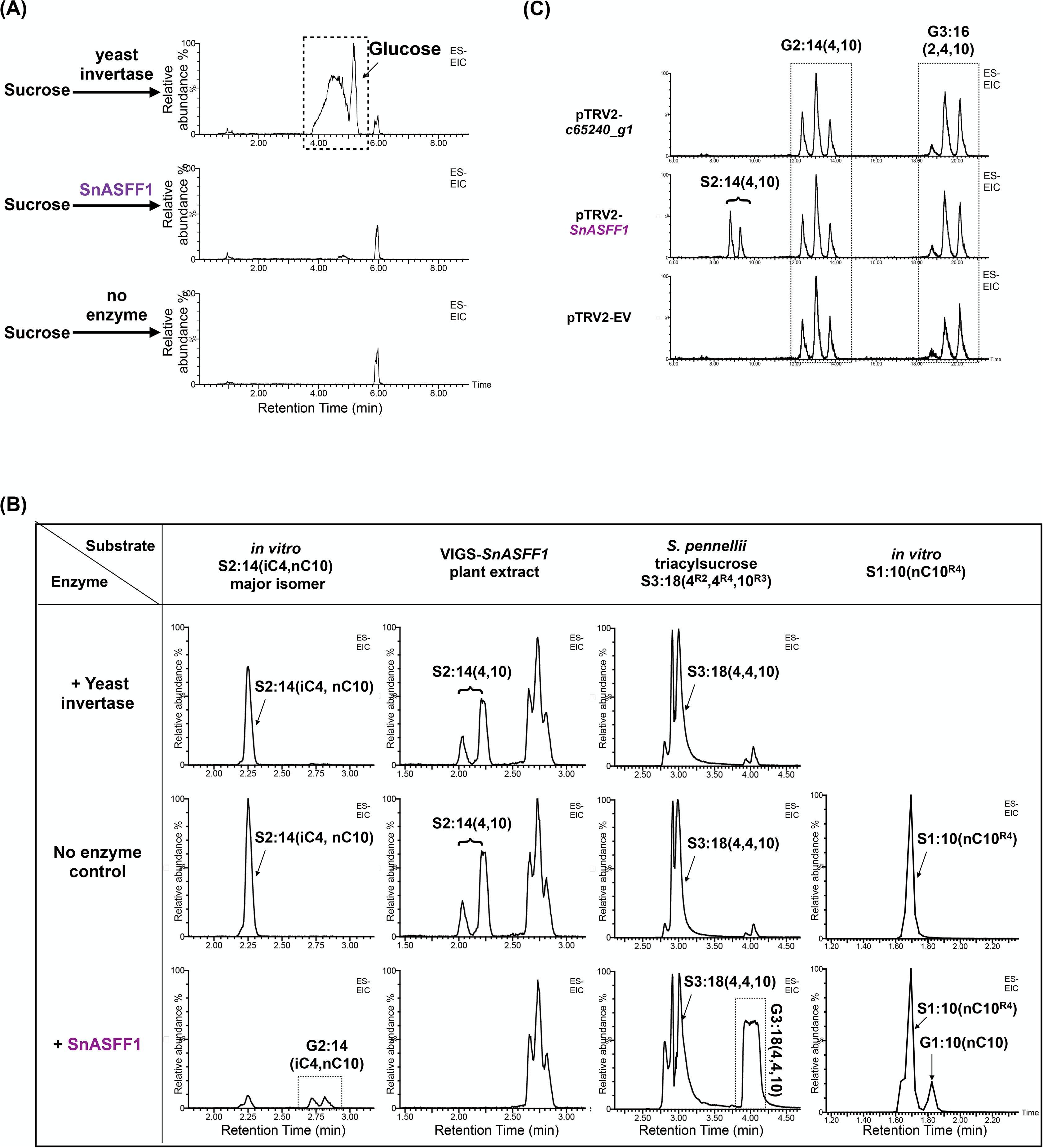
SnASFF1 cleaves mono-, di- and triacylated sucroses but not unmodified sucrose while yeast invertase cleaves unmodified sucrose but not acylsucroses. **(A)** LC-MS analysis of *in vitro* assays with unacylated sucrose showing abundant glucose accumulating in enzyme assay with yeast invertase (top) but not with SnASFF1 (middle). Extracted ion chromatogram (ESI-) showing glucose products (*m/z* 179.06). **(B)** Representative LC-MS of *SnASFF1*-targeted, *c65240_g1*-targeted and empty vector VIGS plants showing diacylsucrose accumulation in VIGS-*SnASFF1* but not in VIGS-*c65240_g1* plants. Combined extracted ion chromatogram showing formate adducts of S2:14 (*m/z* 611.29), G2:14 (*m/z* 445.26) and G3:16 (*m/z* 491.25). **(B)** ESI- mode LC-MS analysis of *in vitro* enzyme assay products indicates that SnASFF1 hydrolyzes *in vitro* and *in vivo* diacylsucrose S2:14(4,10), R_4_-acylated monoacylsucrose S1:10 and R_2_, R_3_, R_4_-acylated triacylsucrose S3:18(4^R2^,4^R4^,10^R3^) purified from *S. pennellii*. Chemical structure of S3:18(4^R2^,4^R4^,10^R3^) was verified by NMR^3^. Combined extracted ion chromatogram showing formate adducts of substrates and products: column 1 and 2, S2:14 (*m/z* 611.29) and G2:14 (*m/z* 445.26), column 3, S3:18 (*m/z* 654.37) and G3:18 (*m/z* 492.31), column 4, S1:10 (*m/z* 541.25) and G1:10 (*m/z* 379.2). Combined extracted ion chromatogram showing formate adducts of S2:14 (*m/z* 611.29), G2:14 (*m/z* 445.26) and G3:16 (*m/z* 491.25).

**Fig. S16.**
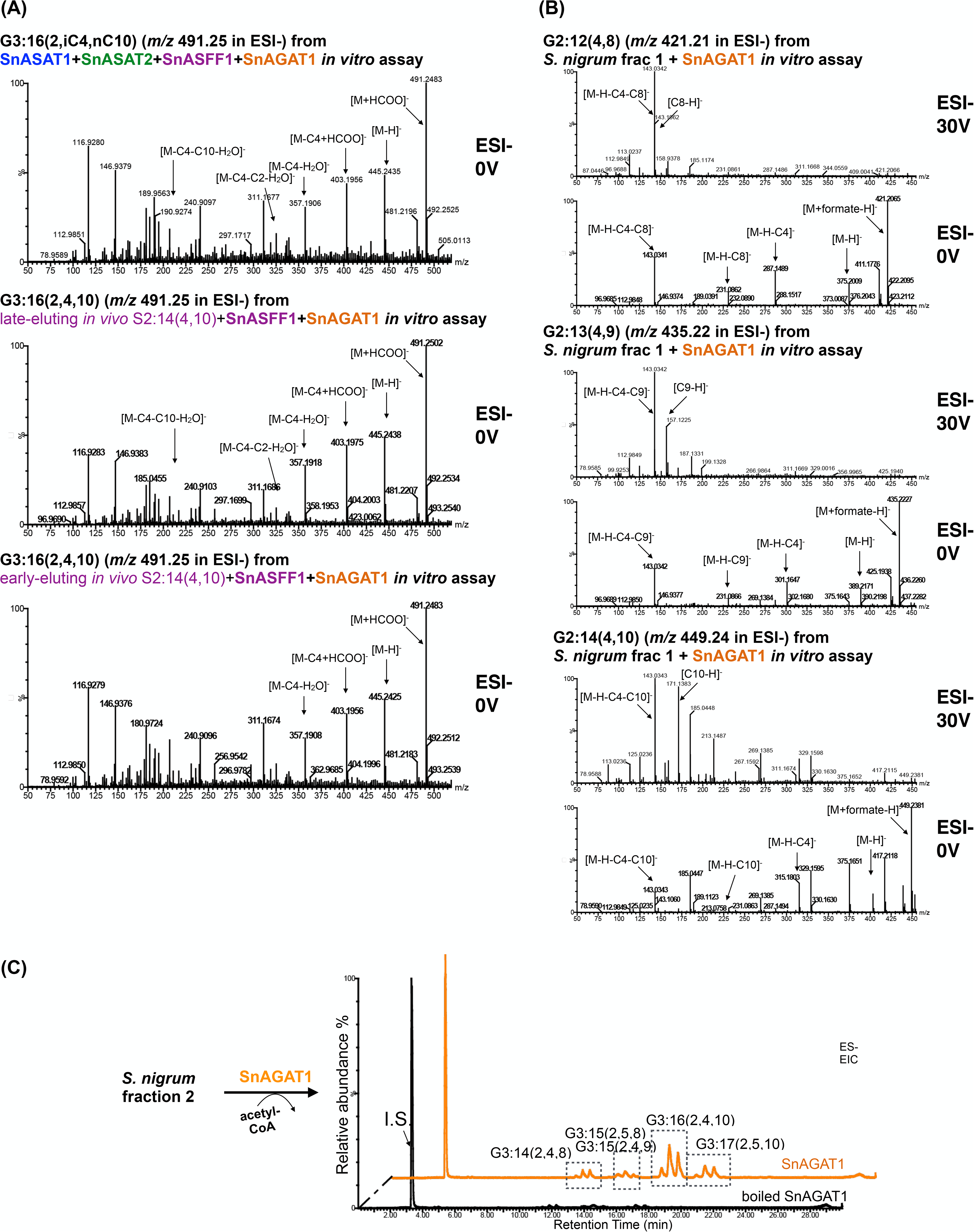
SnAGAT1 forward and reverse enzyme assay products agrees with the acetyltransferase converting diacylglucoses to triacylglucoses. **(A)** The G3:16 isomers produced from SnASFF1 + SnAGAT1 with *in vitro* (top) and *in vivo* (bottom two) S2:14 isomers all generate mass spectra (0V) that are comparable to *S. nigrum* G3:16 (Fig. S1C). Fragmentation of these G3:16 in 0V ESI- mode are characterized by the loss of C4, C10 and C2 ketenes. **(B)** Mass spectra of G2:12, G2:13 and G2:14 diacylglucoses produced from SnAGAT1 reverse enzyme assay with *S. nigrum* extract fraction 1 all agrees with losing an acetyl group. **(C)** LC-MS analysis of enzyme assay products from SnAGAT1 activity (orange trace) shows SnAGAT1 acetylating *S. nigrum* fraction 2 diacylglucoses in the presence of acetyl-CoA. Combined extracted ion chromatogram (EIC) under negative electrospray ionization (ESI-) showing telmisartan as internal standard (I.S.) (*m/z* 513.23) and formic adducts of G3:14, G3:15, G3:16 and G3:17. The corresponding *m/z’s* of these compounds are listed in Table S1 under otherwise specified.

**Fig. S17.**
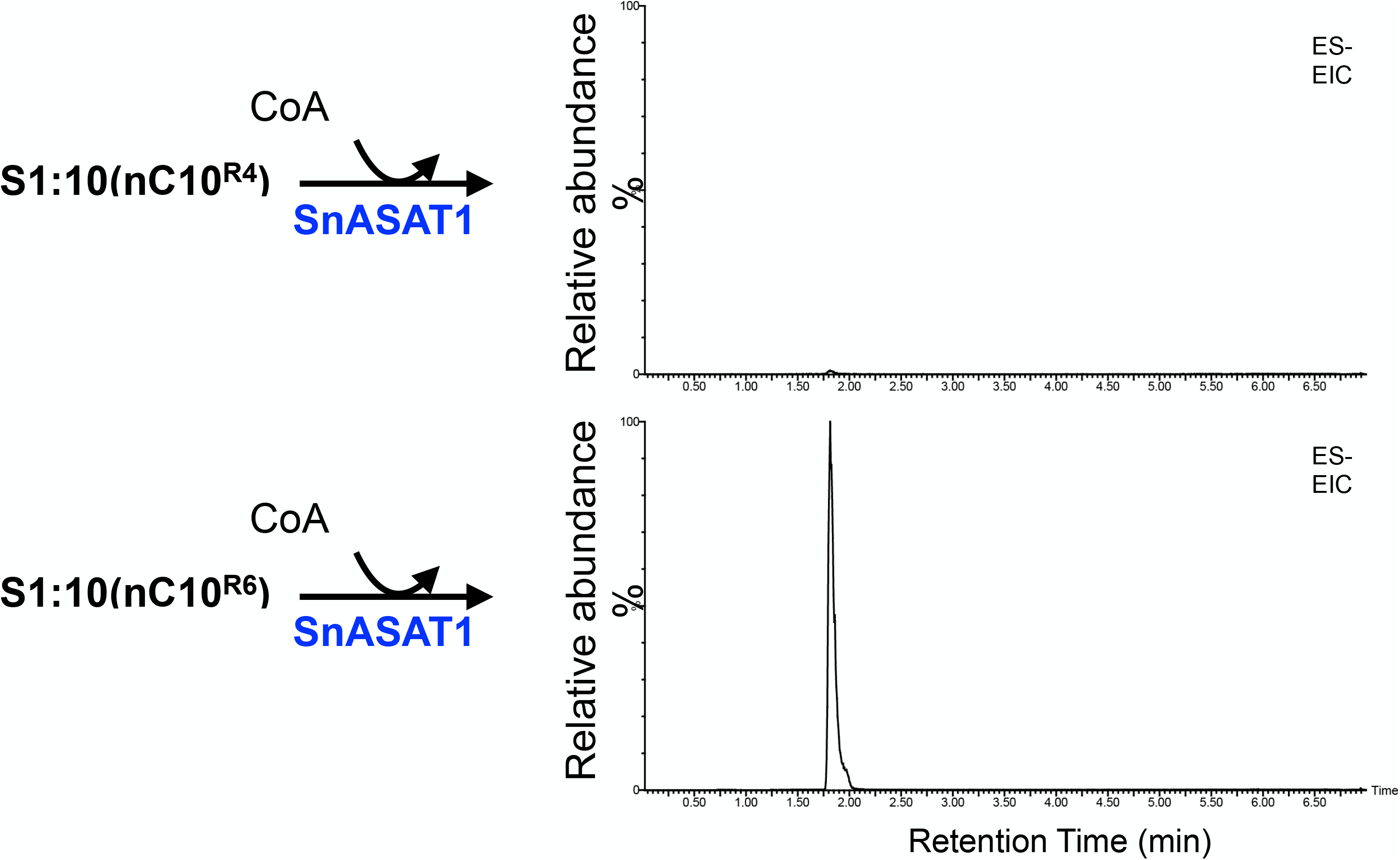
SnASAT1 reverse activity deacylates S1:10(10^R4^) but not the rearranged S1:10(10^R6^) isomer. LC-MS analysis of *in vitro* enzyme assays with purified S1:10(10^R4^) and S1:10(10^R6^) showing SnASAT1 reverse activity diminishes S1:10(10^R4^) accumulation (top) while S1:10(10^R6^) accumulation (bottom) is unaffected. Extracted ion chromatogram showing formate adducts of S1:10 (*m/z* 611.29).

**Fig. S18.**
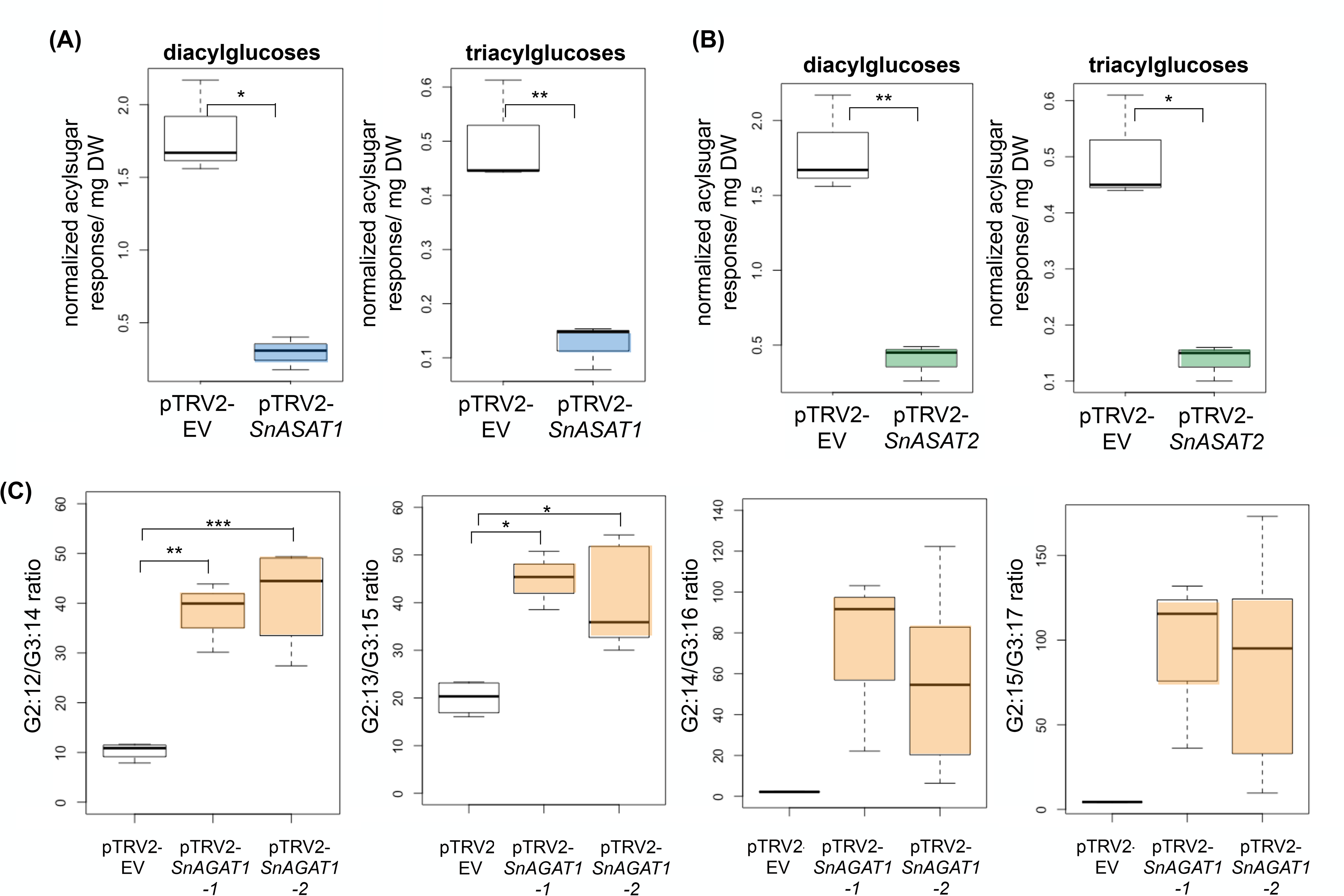
Independent VIGS validation of the BAHD enzymes on *S. nigrum* acylglucose biosynthetic pathway. Comparison of acylglucose accumulation in **(A)** *SnASAT1*-targeted, **(B)** *SnASAT2*-targeted and empty vector VIGS plants. Independent experiments using different constructs from data shown in Fig. 8. **(C)** Comparison of the ratio of G2:12/G3:14, G2:13/G3:15, G2:14/G3:16, and G2:15/G3:17 in *SnAGAT1*-targeted and empty vector VIGS plants. Two constructs targeting different region on *SnAGAT1* were used. Data for pTRV2-*SnAGAT1-1* shown here are biological replicates of data presented in Fig. 8. Acylsugars were analyzed using LC-MS in ESI- mode. Diacylglucose quantities were measured by integrating peak areas of G2:12, G2:13, G2:14 and G2:15, whereas triacylglucose quantities were measured by integrating peak areas of G3:14, G3:15, G3:16 and G3:17. The integrated peak areas were normalized to the internal standard telmisartan (for all) and dry leaf weights (for *SnASAT1-* experiment and *SnASAT2*-silencing experiments). The ratio between tri- and diacylglucoses were independently calculated. The corresponding *m/z* of analyzed acylsugars are listed in Table S1 unless otherwise specified. Significant levels are shown (*, *p* < 0.05; **, *p* < 0.01; ***, *p* < 0.001; Welch’s two sample t-test).

